# Thick filament molecular interfaces play a critical role in pathogenesis of hypertrophic and dilated cardiomyopathy

**DOI:** 10.1101/2025.10.03.680256

**Authors:** Debabrata Dutta, Yuri Kim, Carolyn Ho, SHaRe Investigators, Jonathan G. Seidman, Christine E. Seidman, Roger Craig, Raúl Padrón

## Abstract

Hypertrophic (HCM) and dilated (DCM) cardiomyopathy variants in genes encoding the myosin heavy chain (*MYH7*), myosin light chains (*MYL2* and *MYL3*), and cardiac myosin binding protein-C (cMyBP-C, *MYBPC3*) lead to cardiac hypertrophy or dilatation, with abnormal contractility, relaxation, and energy consumption. Here we defined the structural consequences of >200 pathogenic and benign missense variants in these genes by mapping variants onto a cryo-EM-based atomic model of the human cardiac thick filament. We identified HCM variants residing in 31 molecular interfaces of the complex thick filament interactome, including the two main interfaces of the myosin interacting-heads motif (IHM), and interfaces involving the myosin heavy chain, essential and regulatory light chains, and cMyBP-C. Pathogenic DCM missense variants are rare, and altered only interfaces involving the myosin IHM and tails. None of the 21 variants classified as benign were within interfaces. We demonstrate earlier disease onset and adverse outcomes in HCM patients with pathogenic variants within versus outside of molecular interfaces, emphasizing their importance in normal thick filament function and improving risk stratification of patients. The dissimilar distribution of DCM and HCM variants could explain the different features of the two phenotypes.

## Introduction

Hypertrophic (HCM) and dilated (DCM) cardiomyopathies frequently arise from distinct pathogenic variants in genes encoding protein components of sarcomeres, the contractile units of muscle cells ^1^. The morphological and hemodynamic phenotypes of these pathogenic variants are strikingly different, causing thickening of heart muscle, increased contractility and reduced relaxation in HCM but increased ventricular volumes with reduced contractility in DCM. Even different HCM variants within one gene are recognized to produce variable disease severity and adverse outcomes ^2^. The mechanisms accounting for the distinct pathophysiology of HCM and DCM and associated clinical manifestations remain poorly understood.

Sarcomeres are composed of thin filaments, comprised of actin, tropomyosin and troponin molecules, and thick filaments, assembled from multiple copies of myosin, titin, and cardiac myosin binding protein C (cMyBP-C). The recently elucidated 6 Å resolution cryo-EM structure of the human cardiac thick filament C-zone, and its associated atomic model (PDB 8G4L; Fig. 1a), demonstrated a complex network of intra-and inter-molecular interfaces. This atomic model now allows mapping the HCM and DCM variants onto the thick filament at near-atomic resolution (Fig. 1b, d), helping to elucidate their molecular structural pathogenesis.

**Figure 1.**
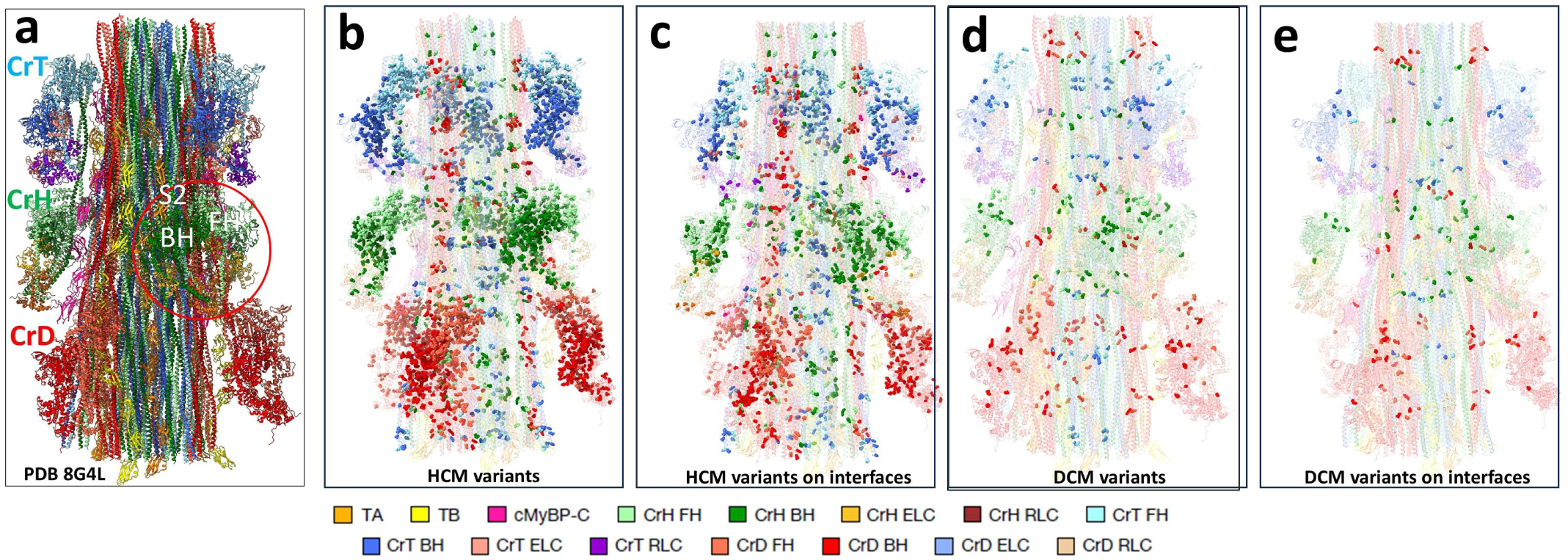
Mapping of HCM and DCM variants on human cardiac thick filament atomic model. **(a)** PDB 8G4L atomic model for 430 Å repeat of the filament C-zone reveals the complexity of structure and numerous interactions in the thick filament ^16^. Three crowns of IHMs are on filament surface (CrH, CrT, CrD) with tails (coiled-coil α-helices), cMyBP-C and titins (TA, TB) in the backbone. Red circle outlines one IHM with blocked (BH) and free (FH) head, and sub-fragment 2 (S2). **(b)** Mapping of the 233 *MYH7*, *MYL2*, *MYL3* and *MYBPC3* pathogenic HCM variants curated from ClinVar onto model in **(a)** (Supplementary Tables 1, 3, 4), color-coded according to protein. **(c)** Same as **(b)** but showing only those variants in molecular interfaces (Supplementary Table 1). **(d)** Mapping of the 25 curated pathogenic DCM *MYH7* variants (Supplementary Tables 1, 2). **(e)** Same as **(d)** but showing only variants in interfaces. See Supplementary Fig. 2 for mapping of benign variants. Abbreviations: TA (titin A), TB (titin B), CrH/T/D (Crowns H/T/D), BH (blocked head), FH (free head), ELC (essential light chain), RLC (regulatory light chain), cMyBP-C (cardiac myosin protein-C).

There are numerous pathogenic *MYH7* missense variants that alter a single amino acid in the cardiac myosin heavy chain (MHC). The vast majority of these variants cause HCM and rarely, DCM. Uncommonly, HCM is caused by *MYL2*, *MYL3*, and *MYBPC3* missense variants encoding respectively, the regulatory light chain (RLC), the essential light chain (ELC), and myosin binding protein-C (cMyBP-C). Initial analyses of the structural basis for dysfunction caused by MHC variants was suggested by their location in distinct regions, including the myosin head (S1) motor domain (MD), actin-myosin interface, nucleotide-binding pocket and converter region, and myosin rod ^3^. These insights suggested how variants impair MD energy transduction but did not account for the prominent relaxation and energetic deficits that characterize HCM and promote adverse outcomes. A crucial subsequent advance was the finding that in relaxed muscle, the two heads of the myosin molecule interact with each other and with the tail ^4 5 6^, forming the myosin interacting-heads motif (IHM) ^7^ (Supplementary Fig. 1a), which establishes an energy-saving, super-relaxed (SRX) state ^8 9^. The structure of the IHM ^5 10 7 11^ provided important new insights. A full mapping of pathogenic HCM and DCM variants onto a quasi-atomic homology model of the human β-cardiac myosin IHM (PDB 5TBY) revealed that myosin variants, while scattered over the heads and tails, were over-represented (i.e. clustered) in interaction sites (interfaces) between the myosin blocked and free heads (BH, FH) and between the BH and sub-fragment 2 (S2) of the IHM ^12^. By impairing IHM formation (BH-S2 interface) and stability (BH-FH interface), more heads would be released and the SRX state compromised, thereby accounting for increased contractility, impaired relaxation, and increased energy consumption that characterize HCM pathogenesis ^12 13 14 15^. While informative of how the structural location of pathogenic variants influenced biophysical properties, these analyses did not provide insights into parameters that contribute to the considerable clinical diversity observed in HCM patients ^2^

The recent cardiac thick filament cryo-EM structure has expanded this restricted scope from a single-IHM to an intact, native thick filament, in which the IHMs are in their natural environment together with cMyBP-C and titin ^16^. This allows analysis of the structural impact of HCM and DCM variants in a native filament. The giant, highly complex 5.9 MDa atomic model of the 430 Å repeating unit of the thick filament C-zone (PDB 8G4L) includes 123 polypeptides, constituting 9 IHMs formed by three different 3-fold symmetric myosin crowns (so-called CrH, CrT and CrD), three different paths of the myosin tails associated with the three types of myosin crown, three cMyBP-Cs, and six 11-domain titin strands, all organized in three radial sectors (Fig. 1a). This intricate structure leads to a complex interactome (Supplementary Fig. 1b), with 32 protein-protein interfaces (regions where two proteins make direct physical contact (<5 Å) in several residue-residue interactions) (Supplementary Table 1), which underlie the functioning of the relaxed thick filament and its SRX state ^16^.

Here, we have used the atomic model to reveal how 258 distinct HCM and DCM pathogenic/likely pathogenic variants (collectively denoted “pathogenic”) curated from ClinVar (see Materials and Methods; Supplementary Tables 2, 3) could differentially disrupt crucial protein-protein interfaces, leading to differently altered cardiac contractility, relaxation, and metabolic activities. Benign and likely benign variants (collectively denoted “benign”), not producing disease, were used as a control (Supplementary Tables 2, 3). We found pathogenic HCM variants on either or both sides of 31 interfaces involving the MHC: between the BH-FH and BH-S2 within the IHM, and MHC with RLC, ELC, cMyBP-C, and myosin tails (Fig. 1c, Supplementary Table 2). No HCM variants were found on the titin A – LMM-T (light meromyosin) and titin-titin interfaces (Supplementary Table 1). As detailed clinical information is not provided in ClinVar, we harnessed the Sarcomeric Human Cardiomyopathy Registry (SHaRe) that links pathogenic variants with longitudinal manifestations of disease and outcomes in HCM patients ^2^. From comparative analyses of the clinical data associated with ≥ 40 patients with a pathogenic variant in the same structural location, we provide qualitative evidence for location-dependent functional consequences.

DCM pathogenic missense variants in thick filament proteins, curated from ClinVar, were only found in interfaces involving the IHM of MHC (n=4, Fig. 1e) and in the S2 and LMM interfaces of the tails (n=10; Fig. 1e, Supplementary Fig. 1a, Supplementary Table 2). The 21 benign variants were only in MHC and cMyBP-C (Supplementary Fig. 2b) and, strikingly, were not in any interface. The dissimilar distributions of HCM and DCM variants provide new insights into their molecular pathogenesis.

## Results

We used the atomic model of the 430 Å repeat of the human cardiac thick filament C-zone (PDB 8G4L; Fig. 1a, Supplementary Fig. 2a) to map the 233 HCM, 25 DCM, and 21 benign variants from the ClinVar database within MHC, ELC, RLC and cMyBP-C (Supplementary Tables 2, 3, Supplementary Datafiles 1, 2, Methods). While the 6 Å model lacks amino acid side-chain densities, it provides excellent overall information on amino acid location, and molecular interfaces are well defined. Interfaces are located where molecules contact closely (<5 Å), establishing electrostatic interactions that are ubiquitously present in the thick filament, and key for assembly and function. We can therefore make qualitative conclusions about the contribution of HCM and DCM variants to disease pathogenesis by assessing their distribution on the thick filament structure and within interfaces. We summarize these structural findings for HCM and DCM variants in comparison to benign variants. Using the considerable clinical data in SHaRe on HCM patients, we considered whether pathogenic variants that perturb molecular interfaces influenced the clinical features and adverse outcomes.

### HCM variants

#### Interfaces involving IHMs

117 HCM variants in *MYH7* alter the heavy chains of the IHMs of CrH, CrT and CrD (Supplementary Fig. 3, Supplementary Table 3). Forty-one of these are in the two internal interfaces of the IHM (BH-FH, BH-S2; Fig. 2, 3) and 21 on IHM interfaces with different tails (Fig. 4, Supplementary Table 1, 4). Of the 41 variants internal to the IHM, 24 are either side of the BH-FH interface (interface “1a”, Supplementary Fig. 1b, Fig. 2) and 17 on either side of the BH-medial S2 interface (“1b”, Supplementary Fig. 1b, Fig. 3). The additional 21 variants involving the IHM are present in interactions of the FH and BH with S2 and LMM tails (“1c” – “1i”, Supplementary Fig. 1b, Supplementary Table 4). The FH participates in four interactions, each impacted by variants: (1) “1d” with a nearby LMM-D (3 variants) (Fig. 4a left), (2) “1c” between its CM loop and its own S2 (4 variants) (Fig. 4b right) or “1h” with tail T distal S2 (5 variants; see Supplementary Fig. 4 for S2 nomenclature) (Fig. 4a right), (3) “1e” with LMM-D (5 variants) (Fig. 4b left). The BH is involved in interaction “1f” with the tail H medial S2 (4 variants) (Fig. 4c). The high number of pathogenic variants in interfaces involving the IHM highlights their importance in normal filament function (see **Discussion**). The remaining 54 motor domain variants, not in interfaces, are likely to be involved in critical myosin ATPase, actin binding, and converter functions ^3^, or may impact interface structure allosterically ^15 17^. Thirty-seven HCM variants are on the coiled-coil α-helices of the S2 region, both where it crosses the open hole of the IHM (proximal S2, not involved in interfaces), and where it interacts with the BH (medial S2), and beyond the IHM (distal S2) (Fig. 3, Supplementary Fig. 4, Supplementary Table 3).

**Figure 2.**
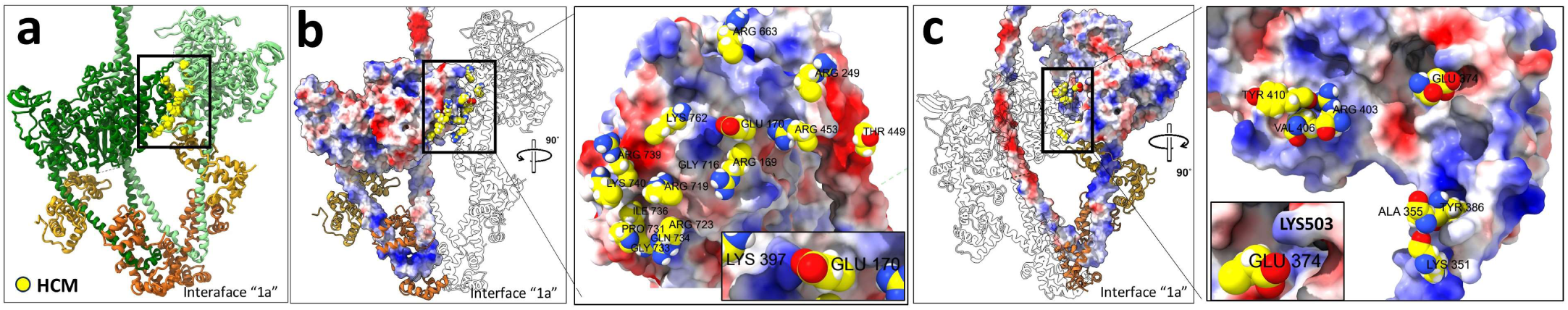
Twenty-four HCM variants in *MYH7* are present in the BH-FH interfaces of CrH and CrT IHMs. **(a)** The interface between the BH (dark green) and the FH (light green) of CrH is shown in the black box. The 24 yellow-highlighted variants in the interface (type “1a”, Supplementary Tables 1, 4) involve seven (4 charge-changing, 3 non-charge-changing) on the BH and 17 (13 charge-changing, 4 non-charge-changing) on the FH. The same occurs on the CrT IHM (not shown). **(b)** Surface charges on the BH are colored blue (positive) and red (negative), and the BH-FH interface (black box) is rotated and zoomed to show FH residues (yellow), opposite in charge to corresponding patches on the BH, demonstrating primarily electrostatic interactions between the two heads. As one example, FH Glu170 interacts with BH Lys397 (inset). The E170K/R charge-reversal variant (Supplementary Table 4) will weaken this interaction. **(c)** Surface charges on the FH are now similarly colored to **(b)**, and the BH-FH interface (black box) rotated and zoomed, showing the BH residues involved (yellow), opposite in charge to the charged surface of the FH, e.g. BH E374 interacts with FH K503 (inset). The charge-loss E374V variant will weaken this interaction.

**Figure 3.**
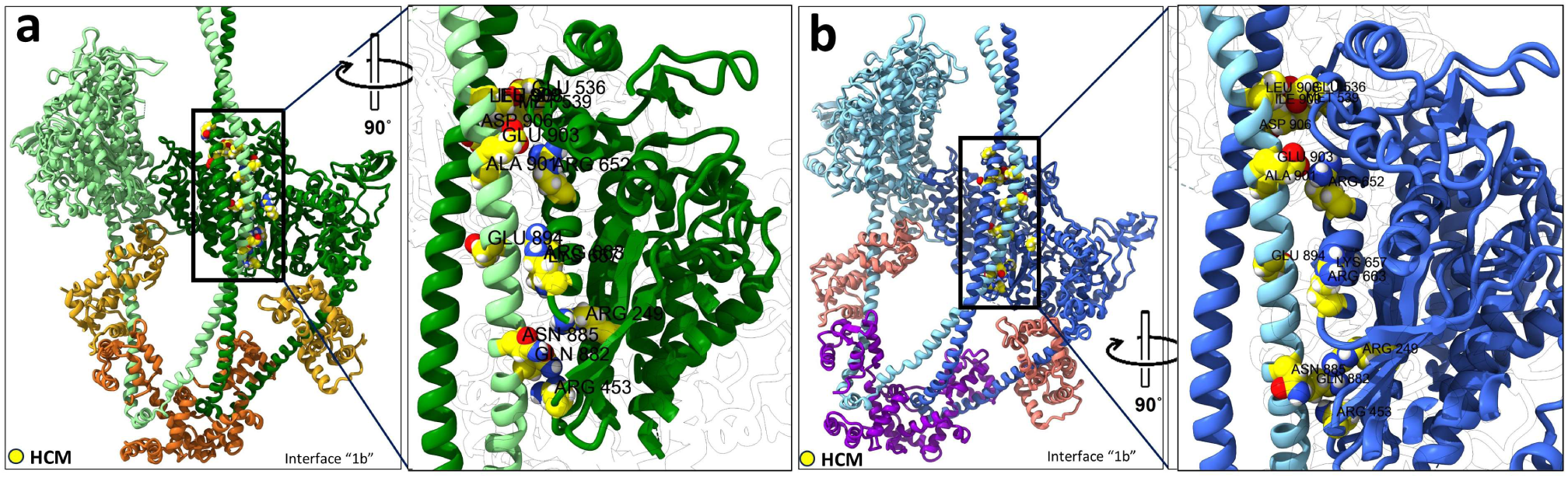
Seventeen HCM variants in *MYH7* occur in the BH-S2 interfaces of CrH and CrT IHMs. **(a)** Interface (type “1b”) between BH and S2 of CrH IHM (black box), rotated and zoomed to show the seventeen residues involved, seven in the BH and ten in S2 (Supplementary Table 4). The charge-changing variants Arg249Gln, Arg453Cys/Ser in this interface are simultaneously present in the BH-FH interface (Fig. 2, Supplementary Fig. 17), producing a dual IHM destabilizing impact. **(b)** Similar interactions occur in the BH-S2 interface of CrT.

**Figure 4.**
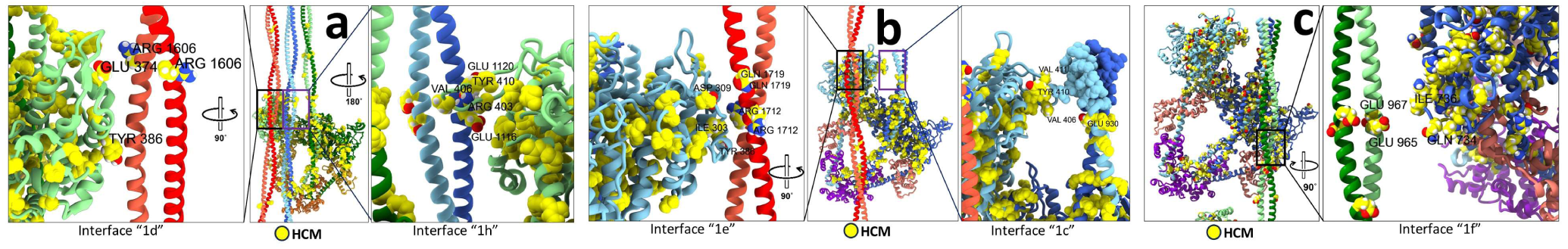
HCM *MYH7* variants are in the IHM BH and FH interfaces with nearby tails. **(a)** Three HCM variants are present in the CrH FH – LMM D interface, two on the FH and one on LMM (type 1d, Supplementary Tables 1, 4). Two are charge-changing, which would impair electrostatic interaction between the FH and LMM-D (zoomed left). On the CrH FH – tail T-distal S2 interface (type 1h), there are five HCM variants, three on the FH side, and two on the distal S2 side (Supplementary Table 4). Several of these, on both sides of the interface, are charge-changing or charge reversal, again impairing interfacial electrostatic interactions (zoomed right). **(b)** There are five variants in the CrT FH – LMM-D interface zoomed on left (type 1e, Supplementary Tables 1, 4). On the CrT FH – tail-T S2 interface, zoomed on right, three HCM variants in the flexible CM loop could interact with Glu930 in the negatively charged Ring 1 of S2 (type “1c”, Supplementary Tables 1, 4) **(c)** On the CrT BH – tail H distal S2 interface (type 1f, Supplementary Tables 1, 4), four HCM variants are present (zoomed on right), two on the BH, and two charge-reversing on tail H distal S2, which could impair the electrostatic interaction between CrH BH and tail H distal S2.

#### Interfaces involving myosin light chains

12 *MYL2* and 5 *MYL3* HCM variants (Supplementary Table 2) are on six interfaces (type “2”, Supplementary Fig. 1b, Supplementary Table 1, 4): (1) BH ELC with MD (“2a”) or lever arm (“2b”, Fig. 5a), (2) FH ELC with MD (” 2a’ “) or lever arm (” 2b’ “, Fig. 5b), (3) BH and FH RLCs (“2c”, Fig. 5c), and (4) BH ELC and S2 (“2d”, Fig. 5d). Notably, *MYH7* variants alter the MHC on the opposite sides of these interfaces (Fig. 5, Supplementary Table 4).

**Figure 5.**
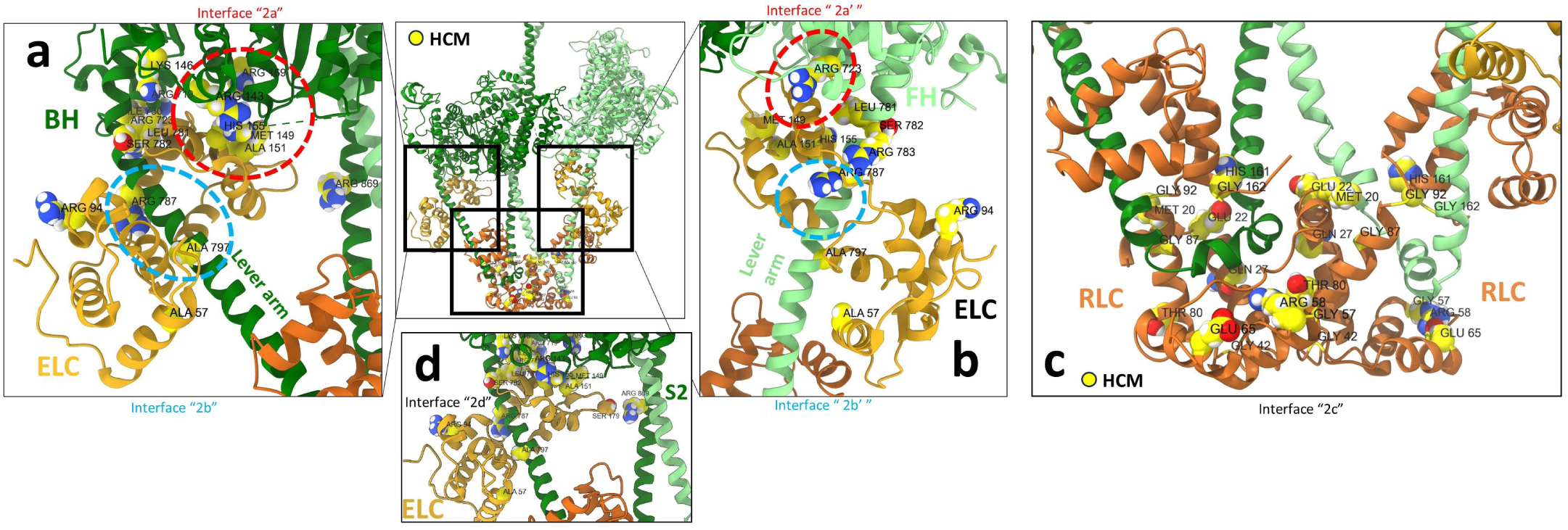
Thirty-two *MYH7*, *MYL2*, and *MYL3* variants occur in six interfaces involving the myosin light chains. **(a)** Eight *MYH7* and three *MYL3* variants are in the BH MD-ELC interface (red ellipse, type 2a, Supplementary Tables 1, 4), which could impair docking of the BH MD on the ELC and thus the specific bent-head conformation of the IHM. Four *MYH7* and one *MYL3* variants are on the BH lever-arm at its ELC interface (blue ellipse, type “2b”, Supplementary Tables 1, 4), which could impair docking of the ELC on the lever arm. **(b)** Similarly, there are three *MYH7* variants on the FH MD at its interface with the FH ELC (red ellipse, type 2a’, Supplementary Tables 1, 4), five *MYH7* and one *MYL3* variant at the FH ELC – lever arm interface (blue ellipse, type 2b’). These variants may have similar impact to the BH variants. **(c)** Three *MYL2* variants on the BH RLC and two on the FH RLC, at the RLC-RLC interface (type 2c, Supplementary Tables 1, 4), may interfere with normal formation and functioning of the IHM or with RLC phosphorylation. The same variants as **(a)-(c)** will occur on the CrT IHMs (not shown), with similar consequences. **(d)** *MYH7* variant Arg869His is in the interaction site 2d (Supplementary Tables 1, 4) between S2 and the BH ELC.

#### Interfaces involving cMyBP-C

16 HCM missense variants (13 *MYH7*, 3 *MYBPC3*) occur in 2 key interfaces (type 3, Supplementary Table 1, Supplementary Fig. 1b), involving: (1) cMyBP-C’s stabilization of IHMs (cMyBP-C domains C5, C8 and C10, 3 variants, interacting with CrH and CrT FHs, 5 variants), and (2) cMyBP-C’s binding to the thick filament through interaction with LMM-Ds (7 variants) (Supplementary Table 1, 4). The interfaces (Fig. 6a) are: (1) C10 with LMM-D and CrT FH (“3d”, Fig. 6b,c); (2) the C9-C10 linker with LMM-D (“3e”, Fig. 6d); (3) C9 with LMM-D (“3c”, Fig. 6e); (4) C8 with CrH FH MD (“3b”, Fig. 6f,g); and (5) the C5 insert with CrH FH RLC (“3a”, Supplementary Fig. 5). All three *MYBPC3* variants were on the surface of domains C5 (Thr750Met), C8 (Arg1022Pro), and C10 (Arg1205Trp/Pro). Of the remaining 16 HCM *MYBPC3* variants, 7 resided on exposed surfaces, not involved in interfaces, while 9 were buried within their respective domains (Supplementary Fig. 6).

**Figure 6.**
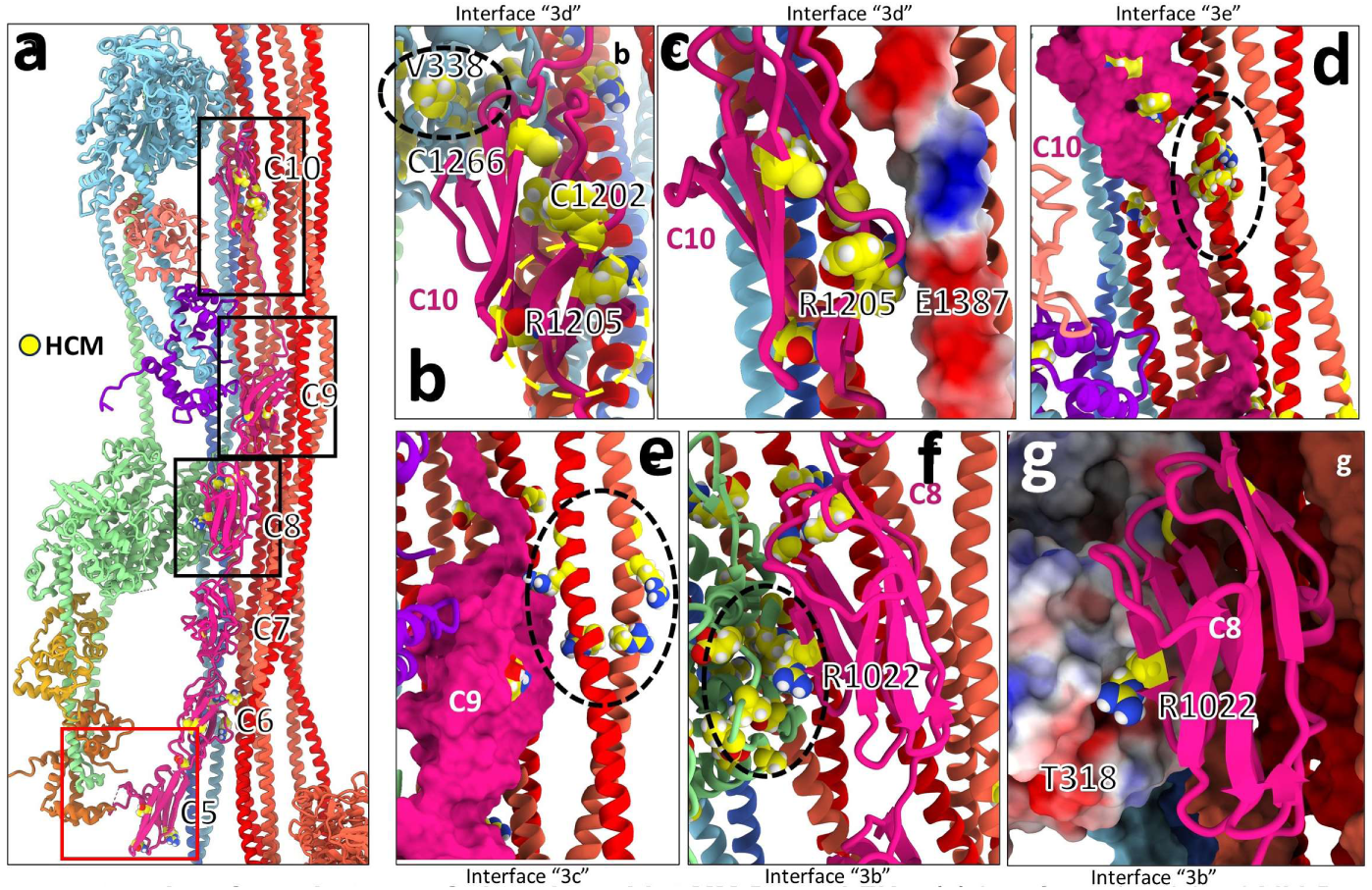
HCM *MYH7* and *MYBPC3* variants are present in interfaces of cMyBP-C with LMM-Ds and FHs. **(a)** Interfaces involving cMyBP-C domains C10, C9, C8, and C5 (boxes) with LMM-Ds (red coiled-coils) and FHs (blue, green). **(b)** Zoom of top box in **(a)** showing interface 3d (Supplementary Tables 1, 4) of C10 with CrH FH MD, containing *MYH7* variant Val338Ala/Met (ellipse). **(c)** Zoom of other part of interface “3d” (yellow circle in **(b)**) between C10 and its electrostatic binding site on LMM-D (blue, positive and red, negative, surface charge representation). Charge-loss mutation of Arg1205 (Supplementary Table 4) would weaken the binding of cMyBP-C to the thick filament, compromising function. **(d)** Variants on interface “3e” of an LMM-D (black ellipse) with C9-C10 linker. **(e)** Zoom of middle box in **(a)** showing variants in LMM-D (black ellipse) at interface 3c with C9 (Supplementary Table 4). **(f)** Zoom of bottom black box in **(a)** showing 4 variants on CrH FH MD at interface 3b with C8 (black ellipse, Supplementary Tables 1, 4), which also interacts with LMM D. **(g)** Zoom of ellipse in **(f)** depicting surface charge in CrH FH MD interacting with positively charged R1022 in C8, which would be weakened with the charge-loss variant R1022P. Bottom red box in **(a)** shows interface “3a” between C5 insert (with variant Thr750Met) and the CrH FH RLC involving *MYL2* HCM variant (Arg58Gln) (Supplementary Fig. 5).

#### Interfaces involving myosin tails

We found 36 HCM *MYH7* variants in 8 interfaces involving tail-tail interaction (Supplementary Fig. 1b type 5, Fig. 7, Supplementary Table 1). Each tail from the three crowns of IHMs is formed by the coiling of two α-helices, one from each MHC, in register (Fig. 1a, Supplementary Fig. 1a). A specific tail variant therefore appears in the same location in pairs (Fig. 7a), with one variant on each α-helix, leading to a total of 6 copies of each variant in the 3-fold symmetric thick filament. In the filament, these 6 copies appear as stripes coming from the T, H, and D tails (Fig. 7e, Supplementary Fig. 11). We found that HCM variants (Supplementary Table 4) occur in three inter-LMM interfaces: between LMM-Ts staggered by 430 Å in the filament core (Supplementary Fig. 1b, “5f”, Fig. 7e), between LMM-Hs in the backbone shell (Supplementary Fig. 1b, “5d”) (not shown), and between LMM-Ds (LMM D sheets, “5h”, Fig. 7b) on the filament surface. HCM variants were also in five interfaces involving LMMs and distal S2s (Supplementary Fig. 7c,d): (1) LMM-H and tail D distal S2 (Supplementary Fig. 1b, “5b”) on the backbone shell (Supplementary Fig. 7c,d), (2) LMM-D and tail T distal S2 (Supplementary Fig. 1b, “5c”) (not shown) on the backbone surface, (3) LMM-D and tail D distal S2 (Supplementary Fig. 1b, “5a”)(Fig. 7b, Supplementary Fig. 9b), (4) LMM-H and tail H/T distal S2s (Supplementary Fig. 1b, “5e”) (Fig. 7c,d), and (5) H and T tail distal S2s (Supplementary Fig. 1b, “5g”, Supplementary Fig. 8a).

**Figure 7.**
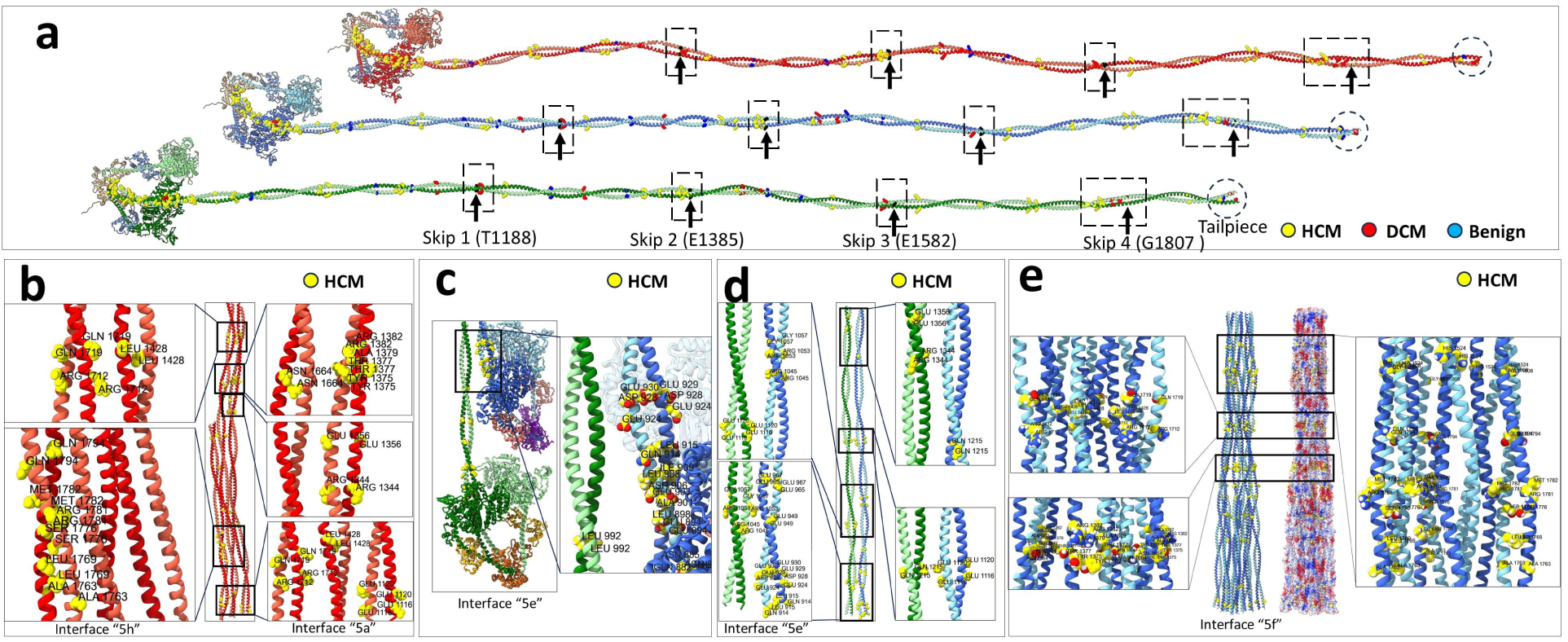
HCM *MYH7* variants are present on LMM-LMM and LMM-S2 interfaces. **(a)** The 58 HCM, 10 DCM and 8 benign variants mapped on tails H, T and D of the human cardiac thick filament C-zone (Supplementary Table 4). These tails (S2+LMM; Supplementary Fig. 1a) are shown in their relative axial positions along the filament backbone. The four skip residues (T1188, E1385, E1582, G1807) are labeled with black vertical arrows. **(b)** LMM-D – tail D distal S2 interfaces (type 5a) assemble LMM sheets on the backbone surface, forming a binding platform for cMyBP-C ^16^. Five groups of variants that could impair formation of these sheets are shown in the black boxes. **(c)** Tail H distal S2 – tail T distal S2 interfaces (type 5g) assemble H and T distal S2s on the outer shell of the backbone; variants Glu924Lys and Asp928Asn are located on tail H distal S2 Ring 2 in this interface. **(d)** Variants are present on tail H and T distal S2 type 5e interface with LMM H (Supplementary Tables 1, 4; center and zoomed). **(e)** There are three groups of variants in LMM-T – LMM-T type 5f interfaces in the filament core (black boxes and zooms), regularly spaced according to their different positioning on the 430 Å repeat. Electrostatic interaction in these interfaces is shown in surface charge depiction at center-right.

#### Interfaces involving titin strands

Two titin strands (TA, TB) position the three myosin tails (H, T, D) to form the thick filament backbone ^16^. We found 22 HCM variants altering the MHC tail at interfaces with titin (Supplementary Tables 1, 4). Sixteen of these variants are on TB interfaces with: (1) LMM-H (Supplementary Fig. 1a, “4a”), (2) tail D distal S2 (“4b”, Fig. 8c), and tail D medial-distal S2 (“4e”, Fig. 8a,b); 6 are on TA interfaces with: (1) LMM H and tail H distal S2 (“4c”, Fig. 8d, f). Tails T are mostly positioned by interactions with tails H and have very few interactions with titin ^16^. Indeed, we found no HCM variants on interface “4d” TA – LMM-T (Supplementary Table 4).

**Figure 8.**
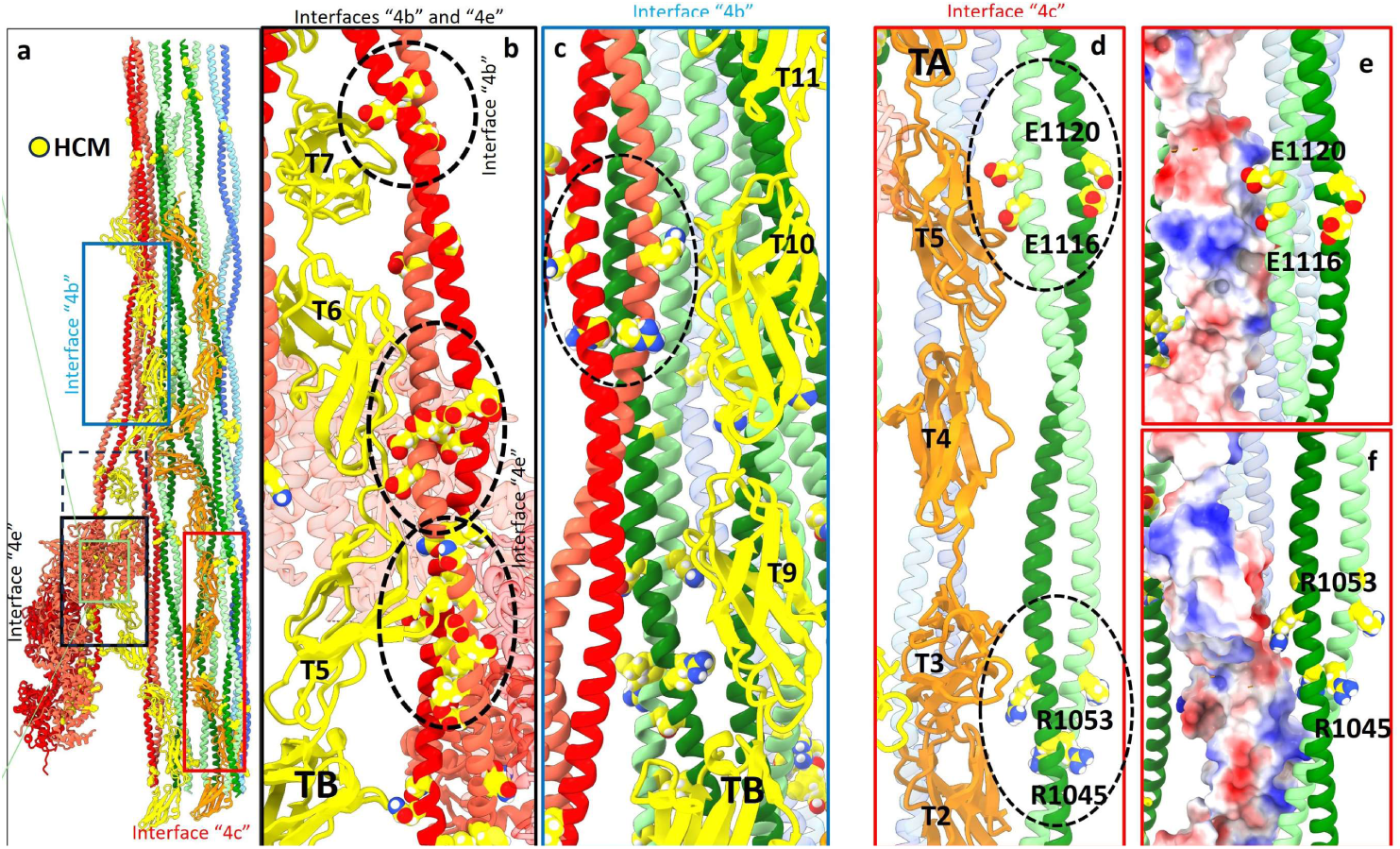
22 HCM *MYH7* variants are present in titin-myosin tail interfaces. **(a)** Overview showing interfaces 4b, 4c and 4e of TA (orange) and TB (yellow) with LMMs and S2 (Supplementary Tables 1, 4; Supplementary Fig. 1b). **(b)** Zoom of black box in **(a)** showing tail D HCM variants at interface 4e with TB domains T5 and T6 (ellipses) and interface 4b with TB domain T7. There are 8 *MYH7* variants on interfaces 4e (Supplementary Tables 1, 4), some in clusters: at TB domain T4 (Gln882Glu, Asn885Lys, not shown), T5 (bottom ellipse: Asp906Gly, Ile909Met, Gln914His), and T6 (top ellipse: Glu924Lys, Asp928Asn, Glu929Lys). There is an interaction (type 4b, dotted circle) between TB domain T7 and LMM D (Glu965Lys, Glu967Lys). **(c)** Zoom of blue box in **(a)** showing interaction type 4b between LMM D and domain T10 of TB (ellipse) containing 3 LMM variants (Supplementary Tables 1, 4). **(d)** Zoom of red box in **(a)** showing interface type 4c of tail H LMM and distal S2 with TA (ellipses, each containing *MYH7* variants). **(e,f)** Zoom of ellipses in **(d)** showing TA surface charges and electrostatic interaction with tail. Only some examples of HCM variants are shown here (see Supplementary Tables 1, 4 for full list).

### DCM variants

There are 25 DCM variants in the curated list, all in *MYH7* (Fig, 1d, Supplementary Tables 2, 3): 15 are on MHC S1 (Supplementary Figs. 3, 12a) and 10 on the tails (Fig. 7a, Supplementary Figs. 8b, 10, 11, Supplementary Table 3). Of those on S1, four (27%) are on IHM interfaces (Supplementary Fig. 12b), while the 10 tail variants are all on tail-titin or tail-tail interfaces. The Clinvar dataset contains no pathogenic DCM missense variants in *MYL2*, *MYL3*, *MYBPC3,* or *TTN* (Supplementary Table 1, Supplementary Fig. 12b).

#### Interfaces involving IHMs

Of the 15 DCM *MYH7* variants on S1, one (R369Q) is on both the BH-FH (“1a”) and CrH FH – LMM-D interfaces, and two (Glu525Lys, Ile533Val) are on the BH-S2 (“1b”) interface (Supplementary Fig. 12b, Supplementary Table 1). The second copy of each of the latter variants is on non-interface sites of the motor domain (Supplementary Fig. 3 vs. 12b). Interestingly, DCM variant Arg904Pro on S2 is adjacent to two HCM variants (Glu903Gly, Asp906Gly), creating an HCM-DCM hotspot (HCM: Glu903Gly, Asp906Gly; DCM: Glu525Lys, Arg904Pro) on the BH-S2 interface “1b” (Supplementary Fig. 13). Eleven of the 15 DCM S1 variants are not on interfaces.

#### Interfaces involving tails

Of the DCM *MYH7* variants in the tail, one is on medial S2, two on distal S2, and 7 on LMM. As with HCM, these variants occur in stripes, but at different locations (Supplementary Fig. 11). Nine DCM variants are in seven tail-tail interfaces “5a-f, h” (Supplementary Figs. 1b, 10, Supplementary Table 4), five of them in multiple different interfaces (Supplementary Table 4); three are in one tail-titin interface involving TB (with LMM-H and tail D S2)(Supplementary Fig. 14a,b), two involving TA (with LMM-H and S2) (Supplementary Fig. 14d), but none on TA - LMM-T interface (“4d”) (Supplementary Table 4) (similar to HCM, see above).

### Benign variants

We mapped the 21 *MYH7* and *MYBPC3* benign variants (Supplementary Table 2, Supplementary Table 3) on 8G4L (Supplementary Fig. 2b). In contrast to the HCM and DCM variants (Fig. 1c, e, Supplementary Table 4), benign variants were not found on any interface (Supplementary Figs. 4,15,16).

### Clinical Associations with HCM variants

The SHaRe registry links pathogenic variants with longitudinal manifestations of disease and outcomes in HCM and DCM patients ^2^ (Supplementary Datafile 3), data that is not provided in ClinVar. Within SHaRe there are 890 HCM patients with 212 unique pathogenic missense variants in *MYH7* and 73 HCM patients with 41 pathogenic missense variants in *MYBPC3*. Among DCM patients in SHaRe (n=65) there were unique pathogenic missense variants in *MYH7* (n= 32) and *MYBPC3* (n=4). To assess whether the molecular locations of missense variants were associated with different burdens of disease, we compared the clinical manifestations between patients with variants within and outside of an interface or structural region (Supplementary DataFile 4). As we required ≥ 40 patients within the same region, the clinical analyses were restricted to MYH7 variants. Specifically, we compared clinical features in patients with interface variants (BH-FH, n=123; and BH-S2, n=172) to patients with non-interface variants (n=433 and 431, respectively) in the MHC head. Similarly, we compared HCM patients (n=330) with variants across the entire MHC head to patients (n=556) with tail variants, and between patients with tail variants (proximal S2, n=114; medial S2, n=47; remaining tail, n=146) to patients with variants across the entire protein.

Within each structural group we assessed patients’ age-specific disease burden by comparing the mean age when HCM was clinically manifest (e.g., primary diagnosis or left ventricular (LV) thickness ≥ 13 mm) and mean age when a significant LV outflow tract obstruction (≥ 50 mmHg) was identified. Additionally, we assessed the mean age for first occurrence of atrial fibrillation, ventricular arrhythmic composite, heart failure composite, stroke, and death (Supplemental Datafiles 3, 4). Not all information was available for every patient.

HCM patients with pathogenic missense variants within the BH-FH interface were diagnosed at approximately 12 (mean age) years younger than patients with variants elsewhere in the MHC head (adj p= 5.0e-9; Hazard Ratio (HR) = 1.92; Supplemental Datafile 4, Figure 9). The BH-FH interface was also associated with significantly younger mean ages (adj p=2.0e-2, each endpoint) for outflow tract obstruction (HR = 2.18), atrial fibrillation (HR =1.82) and ventricular arrhythmia composite (HR =5.06). The mean ages were approximately 10 years younger for stroke (adj p=2.0e-3, HF=6.36) and heart failure composite (adj p=7.0e-6; HR =2.64).

**Figure 9.**
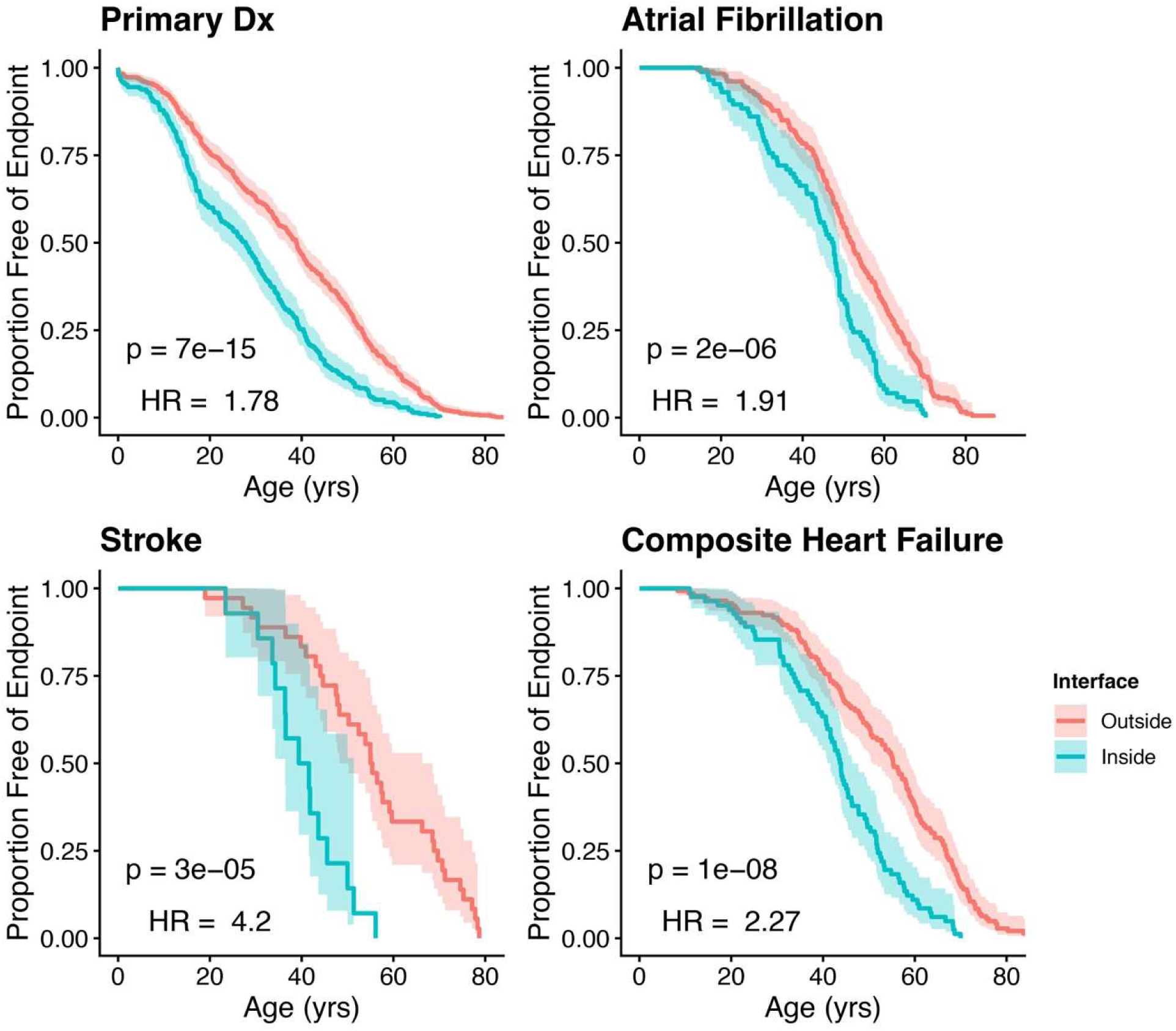
Comparisons of the age of first Primary diagnosis (Primary Dx; top left), atrial fibrillation (top right), stroke (bottom left) and heart failure (bottom right) of individuals with a variant in the BH-FH interface vs those with variants elsewhere in the MYH7 head. Kaplan-Meier curves were calculated for SHARE subjects with (“YES”) an MYH7 variant in the BH-FH interface (n=123, 14, 41, 42, for HCM onset, AF, HF and Death, respectively) and with a MYH7 variant in the head region but not in the BH-FH interface (n=740, 128, 183, 225, HCM onset, AF, HF and death, respectively). Nominal p-values were calculated by the log-rank test (see Methods).

Parallel analyses of variants in the BH-S2 interface to variants elsewhere in the MHC head indicated a modest impact on disease burden (mean ages at diagnosis, adj p =2.0e-2; HR =1.36), heart failure composite (adj p = 0.01, HR= 1.8) and death (adj p = 0.05 e-3, HR =2.44). As this comparison included the influence of variants in the BH-FH interface, we also assessed the disease burden for the BH-S2 interface to all non-interface MHC variants. These analyses indicated that variants in the BH-S2 interface significantly impacted disease (e.g., younger mean ages at diagnosis, adj p = 1.0e-7, HR= 1.68; atrial fibrillation, adj p = 2.0e-4, HR= 1.96; heart failure composite, adj p=5.0e-5; HR =2.2; death, adj p=2.0e-4; HR =3.3), albeit less so than the BH-FH variants.

Notably, clinical data from patients with a variant anywhere in the MHC head compared to the tail showed no significant mean age differences for all endpoints. Similarly, comparisons of variants within each tail segment and the entire protein indicated no substantial increase in disease burden. Notably, the proximal S2 region (32 residues), contains a significant clustering of ClinVar variants (n=11, (p=8.0e-8, Supplemental Table 5). However, among 12 S2 proximal variants in 114 SHaRe patients compared to all patients with other variants, the mean ages were significantly <u>highe</u>r at diagnosis (adj p = 5.0e-5, HR =0.62) and not different for other endpoints. These observations implied that S2 variants are most deleterious when these disrupt protein-protein interactions within the BH-S2 interface.

Taken together these data indicate that *MYH7* variants within the BH-FH and BH-S2 interfaces evoke unfavorable clinical manifestations of HCM (Figure 9). Comparisons of variants within both interfaces compared to all non-interface variants indicate younger mean age of approximately one decade at diagnosis (adj p = 7.0e-15, HR =1.78), onset of atrial fibrillation (adj p = 5.0e-5, HR =1.91), stroke (adj p = 7.0e-4, HR =4.2), composite heart failure (adj p = 1.0e-8, HR =2.27), and death (adj p = 5.0e-4, HR =2.94). We suggest these clinical data reflect that interface variants cause substantial dysfunction within thick filament proteins. In response, the heart initiates early adverse remodeling that accelerates clinical diagnosis. The persistence of dysfunction combined with cardiac remodeling propels earlier onset of adverse clinical events, including atrial fibrillation, stroke, heart failure and death.

## Discussion

The lack of structural information on the human cardiac thick filament has limited our understanding of the molecular pathogenesis of HCM and DCM caused by variants in thick filament proteins. Previous analysis of variants was based on the crystal structure of S1 (2MYS)^3^ and on atomic models of the isolated IHM, and a low-resolution thick filament model based on negative stain EM (5TBY, ^12^; 8ACT, ^18^). The cryo-EM thick filament model (8G4L; ^16^ now places the IHMs in their native context, along with tails, cMyBP-C and titin, revealing 32 molecular interfaces, 30 of which contain pathogenic variants (Supplementary Table 1). We show that the structural impact of pathogenic HCM and DCM variants, and benign variants depends on their different locations in these interfaces. HCM pathogenic variants, curated by ClinVar, occur in four genes (*MYH7*, *MYL2*, *MYL3*, and *MYBPC3*, but not *TTN*) and are present in all 30 interfaces; DCM variants in *MYH7* and are restricted to BH-FH, BH-S2, and tail-tail and tail-titin interfaces. Strikingly, benign variants are not in any interface. These results highlight the fundamental importance of intra- and inter-molecular interfaces, and thus of a properly formed thick filament structure, to normal function. Disruption of these interfaces by HCM or DCM variants may contribute to the abnormal contractility, relaxation, and energy consumption that characterize these diseases.

### Structure-based insights into HCM and DCM

Pathogenic HCM and DCM missense variants could impact thick filament function in two main ways, depending on their location: compromising myosin motor activity or disturbing thick filament structure/stability. Variants in the myosin heads (MHC, ELC, RLC) and proximal and medial S2, which together form the IHM, would directly impact myosin motor function, regulation of SRX, and head availability for actin interactions. Backbone (tail or cMyBP-C) variants could indirectly impact motor function—by disruption of gross filament structure, or disturbance of the delicate interactions that stabilize the IHMs in their off-state (SRX) ^19^. Reduced cMyBP-C levels, the consequence of almost all HCM pathogenic variants in *MYBPC3*, similarly compromise IHM stability ^20^. The larger number of variants in the IHM than in the backbone (Supplementary Table 4) suggests that cardiomyopathy occurs more frequently through direct impact on myosin function. Importantly, the different number of myosin, cMyBP-C, and titin molecules in the C-zone (MHC and light chains, 162 copies; cMyBP-C, 27 copies; titin, 6 copies) means that the impact of a variant in *MYH7*, *MYL2*, or *MYL3* will be potentiated compared with variants in *MYBPC3* or *TTN*. The relative muted effects by a deleterious *TTN* missense variant and the potential for compensation by vast numbers of prevalent *TTN* missense variants found in all genomes may account for the insufficient fulfillment of pathogenic criteria by *TTN* missense variants in the C-zone ^21^.

#### Clustering of variants in molecular interfaces

Within the IHMs and backbone, many of the pathogenic variants are strongly over-represented in the interfaces between or within molecules. The 24 HCM variants in the BH-FH interface (Supplementary Table 4) represent 39% of the total 61 BH-FH interface residues, a 2.8-fold enrichment over the expected number if the 117 HCM head variants were distributed evenly over the head (p=3e-6; Supplementary Table 5). There is also a striking 1.9-fold enrichment in the BH-S2 interface (p=0.009), mainly on the medial S2 side (4.8-fold enriched compared with the tail as a whole; p=5e-6). This clustering of HCM pathogenic variants highlights the central role of the IHM in normal cardiac function: disturbance of the BH-FH or BH-S2 interfaces would be expected to loosen myosin heads and disrupt the SRX state, leading to increased contractility, impaired relaxation, and increased energy consumption that characterize HCM ^12 13 15^. The clinical import of variants in these IHM interfaces is further emphasized by the earlier onset of HCM and increased adverse outcomes reported by the SHaRe registry on HCM patients with these variants (Supplementary Datafiles 1, 4).

ELC and RLC variants that cause HCM are rare ^22^. HCM variants in the interfaces of the ELC with the lever arm (Fig. 5b; Supplementary Table 4) and the motor domain (Fig. 5a, b; Supplementary Table 4), may impair the bent conformation of the BH and FH heads, and the lever arm flexibility required to form a stable IHM through BH-FH and BH-S2 docking. Variants in the BH RLC in its interface with the FH RLC (Fig. 5c) would impact on the fulcrum where the heads connect to S2, and thus flexibility of the head-tail junction, and possibly the RLC phosphorylation mechanism. In contrast to the IHM, relatively few pathogenic variants (35 of 175; Supplementary Tables 3, 5) are in the myosin tail (i.e. distal S2 and LMM), given the length of this region (1018 residues). Strikingly, however, 31 (almost 90%) are in interfaces—in fact multiple, different interfaces—with other tails, heads, cMyBP-C and titin (Supplementary Table 4), highlighting the key role of intermolecular backbone interactions in normal filament assembly and function.

The atomic model 8ACT of a recombinant human myosin IHM construct containing 15-heptads of tail ^18^ confirmed at 3.6 Å resolution that the intra-IHM interfaces BH-FH (1a), BH-S2 (1b), ELC (2a, 2a’, 2d) and RLC (2c) are hotspots of HCM mutations that were previously proposed to cause hypercontractility by destabilizing the IHM ^12^. The 8ACT model shows strong similarity with CrH and CrT IHMs in 8G4L ^16^, allowing further understanding of these interfaces. However, we do not find support for the proposal that the restrictive cardiomyopathy (RCM) ELC E143K variant and the HCM D166V and R58Q RLC variants correspond to BH ELC and FH RLC residues that interact with the core of the thick filament ^18^. The cryo-EM thick filament atomic model ^16^ does not demonstrate these interfaces, which were suggested originally by a (dried and collapsed) negative stain structure ^23^ cf. ^24^. This observation underscores the importance of using the near-native thick filament structure in assessing structural effects of thick filament variants.

The presence of pathogenic variants on both sides of many interfaces (FH/BH, BH/S2, ELC/MHC (Fig. 5), cMyBP-C/myosin head) emphasizes their prominent physiologic functions ^12 14^. In contrast, in titin-tail interfaces, pathogenic variants are identified only in the MHC tail and not in titin.

#### Variant effects depend on the environment

Owing to the asymmetric structure of the IHM, a variant in the myosin head will occur in different environments depending on whether it is in the BH or FH. For example, Glu525Lys in the MD of the BH is present in the BH-S2 interface, which would affect IHM stability. The same variant in the FH is not in any interface and may be inconsequential or affect myosin ATPase or actin binding of the FH. Rarely, a variant occurs in both interfaces of the IHM: HCM variants Arg249Gln and Arg453Cys/His/Ser residing in the FH impact the BH-FH interface, but when on the BH they impact the interface with S2 (Supplementary Fig. 17). Similarly, the DCM variant R369Q within the BH interferes with the BH-FH interface (Supplementary Fig. 12b), and when on the FH, alters interactions with the tail of a neighboring myosin molecule (Supplementary Table 4). Similar considerations apply to the light chains, which have different interactions in the BH and FH. Further complexity arises because the IHMs themselves (CrH, CrT and CrD) are not identical, with different stabilities and interactions in the three locations within a triplet ^16^. This is apparent in the different interfaces of the BH or FH of CrH, compared with CrT, with nearby L, T or D tails (Fig. 4, Supplementary Table 4). There are consequently six different head environments with six possible outcomes from a single point mutation. Thus, variants in R403 in the CM loop may impair FH interaction with its own S2 in CrT and with a different tail in CrH ^16^. The dimeric structure of myosin adds even more complexity, as variant and WT polypeptide chains in a heterozygous variant may assemble to form homo- or hetero-dimers, or a mixture of the two, and these could assemble into fully mutant, fully WT, or mixed thick filaments.

Tail variants also occur in different environments. The tails are coiled-coils composed of in-register α-helices and a variant in any one amino acid could thus affect one α-helix or the other (or both), impacting interactions on the two sides of the tail differently. And the tails from the three different crowns are themselves in different environments in the filament backbone, either forming the filament core (CrT), the filament shell (CrH), or the filament surface, where CrD tails also interact extensively with cMyBP-C ^16^. Thus, a variant in an H, T, or D tail affecting a specific tail-tail or tail-titin interface could have a quite different effect in another tail type, depending on its own, different interactions (Supplementary Fig. 18). We suggest that these considerable structural complexities, in addition to other genetic and hemodynamic differences, contribute to the variable clinical manifestations and outcomes observed in affected family members, despite sharing an identical pathogenic variant ^25^.

#### Most variants involve charge change

The majority of MHC head, tail and cMyBP-C missense variants involve a loss, reversal, or sometimes a gain, of charge, altering the strength of electrostatic interaction in an interface. Most of the 40 HCM (Fig. 2, 3) and 4 DCM (Supplementary Fig. 12b) variants in the BH-FH and BH-S2 interfaces are charge-changing, frequently involving arginine, lysine, histidine, glutamic acid and aspartic acid (Supplementary Table 4). We suggested these destabilize/ stabilize the IHMs, leading to the hyper- or hypo-contractility characteristic of HCM and DCM respectively ^12^. Similar charge changing variants are also in the remaining interfaces (Supplementary Table 4), which may similarly destabilize/stabilize these interfaces.

#### Variants in cMyBP-C interfaces affect IHMs

Pathogenic HCM missense variants were found in two types of interfaces involving cMyBP-C: where it interacts with IHMs and where it docks on the thick filament by binding to a sheet of 3 CrD myosin tails ^16^. Interactions of cMyBP-C domains C5, C8 and C10 with the free heads of CrT and CrH are expected to stabilize the IHM, enhancing the SRX state ^26 27^. cMyBP-C variants that weaken these interfaces on one side, or MHC or RLC variants on the other, would compromise IHM stability, freeing more heads for actin interactions, contributing to the hyperdynamic contractile phenotype of HCM. Disruption of cMyBP-C binding to the filament surface, by variants on either side of the interface of domains C8-C10 (Supplementary Tables 1, 4) with the binding platform of CrD tails ^16^, would be expected to have a similar effect, by reducing or eliminating cMyBP-C’s stabilization of CrT and CrH IHMs. Only variants on the surface of cMyBP-C domains are directly involved in these interfaces with FHs and CrD tails. Analysis of other cMyBP-C variants buried within domains (Supplementary Fig. 6) awaits more detailed structures.

#### Tail variants impair thick filament structure

The complex structure of the thick filament backbone results from precise interactions of myosin tails with each other and with titin ^16^. Our analysis (Fig. 7a) shows that a single tail variant will likely have different structural effects in the three types of tail (T, H, D) due to their strikingly different conformations and environments: these tails interact through eight tail-tail interfaces (“5a-h”, Supplementary Figs. 1b, 9, Supplementary Tables 1, 4), creating the backbone core (LMM-T), a surrounding shell (LMM-H), and the backbone surface (LMM-D). Variants in these interfaces could impair formation of the native backbone structure, disrupting the correct interactions of IHMs with the filament surface (Fig. 4), and thus reducing SRX ^19^. Similar disturbance of filament assembly has been found with skeletal muscle tail variants (Supplementary Fig. 9a,c), some that cause myosin storage myopathy (MSM; ^28^). Notably, an HCM/DCM hotspot near the terminus of the tail, in a region involved in multiple tail-tail interactions ^16^, could impact tail packing, affecting filament formation and stability (Fig. 7a).

Myosin tail variants also occur at interfaces with titin A or B (Supplementary Table 4; Fig. 8, Supplementary Fig. 14): these may also impair filament assembly, which depends on specific interactions at these sites. A striking example is the interface of titin B with negatively charged Rings 1 and 2 on the medial S2 of CrD, where it interacts with the BH. Kinking of TB in this region raises domains TB4-TB6 above the backbone and similarly raises the CrD IHM (65 Å; Fig. 8a black box) ^16^. This interface (“4e”) is unique and not present on the ordered CrH and CrT IHMs, which are close to the backbone. Eight HCM and two DCM variants occur on Rings 1 and 2 where they interact with TB (Fig. 8b). These could unlock the positioning of the CrD IHM above the backbone, altering the functioning of its proposed sentinel/swaying heads ^29 30 16^.

Interestingly, while most MHC tail variants are charge-changing—and would affect filament structure by altering electrostatic interactions with other tails or with titin—nine do not alter charge (Supplementary Table 4). These may alter the local conformation of the coiled-coil, impact tail interactions, or alter IHM structure and SRX by transmission along the coiled-coil (see below).

#### Non-interface pathogenic variants

Variants in regions of myosin and cMyBP-C outside of interfaces may impact function by allosteric influence on interface stability ^15 17^, or by directly affecting myosin enzymatic function or actin-binding capability ^3^. HCM variants in MHC are clustered in the proximal S2 region of the tail, immediately after it leaves the head-tail junction (Supplementary Fig. 4b, Supplementary Table 5) and may impair IHM formation (cf. ^31 32^).

Residues mutated to proline, which disrupts α-helical secondary structure, occur only in the IHM (heads and proximal/medial S2) and never in the distal S2 or LMM, that primarily form the tail (Supplementary Fig. 19). Mutated MHC prolines in the IHMs could still form thick filaments (as the IHM does not have a major role in filament formation) but may have impaired enzymatic or SRX function. In contrast, the distal S2/LMM part of the tail is central to filament assembly, and disruption of its α-helical structure may prevent proper filament formation. Proline tail variants may therefore be lethal explaining why these are never observed.

#### Comparing HCM and DCM

There are ten times more missense variants that cause HCM than DCM, and these altered MHC, light chains, and cMyBP-C, affecting 31 thick filament interfaces. DCM missense variants were in MHC and affected only IHM and tail interfaces (Supplementary Table 4). The reciprocal contractile functions, increased in HCM and decreased in DCM, suggest increased/decreased myosin activities. Charge-changing HCM variants in IHM interfaces reduce charge-charge attraction and disrupt IHM stability ^14 15^, thereby increasing the availability of heads for actin interactions, biophysical changes consistent with increased contractility. Rare variants could potentially increase charge-charge attraction and thereby enhance IHM stability and SRX. For example, DCM Glu525Lys ^33^ may increase attraction between the BH (Lys525) and negatively charged residues on Ring 1 of S2 ^10^ (Supplementary Fig. 13), which could augment stabilization of the BH-S2 interface and thus the IHM, and thereby limit heads available for actin interactions, increasing SRX, and producing hypo-contractility characteristic of DCM ^33^.

While the protein structural differences between variants can be quite subtle, their consequences can be considerable, as illustrated by HCM variant Arg783Cys/His and DCM variant Arg783Pro (Supplementary Table 4, Supplementary Fig. 12). Two contiguous variants in a single interface can do the same: GlyE903Gly and Asp906Gly cause HCM, and Glu904Cys/His causes DCM. This interface also contains DCM variants at Glu525 and Ile533, creating a DCM/HCM hotspot (Supplementary Fig. 13). The consequence of each specific variant may depend on the stabilization (DCM) or destabilization (HCM) of the IHM, through changes in the local interactions of these residues, impact of charge changes, and other mechanisms that may be revealed by regional high-resolution structures. Subtle structural effects may also account for why some MHC tail variants result in HCM and others in DCM. We suggest HCM variants may allow relatively normal filament assembly but disrupt IHM-backbone interactions (Fig. 4) and IHM stability sufficiently to increase head availability for actin interactions, causing hypercontractility. By contrast, DCM tail variants may cause greater disruption of filament structure, perhaps reducing the effective number of functional heads, leading to reduced force transmission and hypocontractility.

### Clinical Utilization of Structural Insights

The uptake of gene-based diagnosis of HCM and strong endorsement by practice guidelines is predicated on the utility for genotype to identify patients at greatest risk for adverse outcomes and to proactively intervene to mitigate these risks ^34 35^. However, the vast numbers of different pathogenic HCM variants that are unique to one or a few families, and the variable severity of manifestations associated with variants in the same gene and between genes has challenged this clinical goal. Longitudinal studies of large genotyped HCM cohorts, such as SHaRe, provide compelling evidence for earlier onset and worse disease in patients with a definitive pathogenic genotype compared to patients with unknown etiologies. The integration of structural data has the potential to substantially refine these insights. By demonstrating that adverse clinical manifestations are linked to MHC variants within interfaces, we identify HCM patients at highest risk for earlier onset of disease, arrhythmias, and heart failure. This information identifies patients who can benefit from careful monitoring and pre-emptive interventions to limit adverse outcomes. By contrast, the accurate identification of HCM patients with a more favorable disease course can reduce surveillance and unnecessary interventions. Further, these data open opportunities to define small molecules that selectively influence interface interactions to normalize function and attenuate disease. We suggest additional high-resolution structures of native thick and thin filaments will advance fundamental insights into sarcomere biology that can improve the care of HCM and DCM patients.

## Conclusion

Genetic variants can in principle occur at any location in the translated protein. At certain, key, functional locations, they may be embryonically lethal and not observed. An example may be the striking absence of pathogenic variants in loop 2 (residues 626-647) of the MHC. This loop is involved in interactions of S2 with both the FH and BH in the IHM and with actin, and as such may be essential to function. At other locations, variants may be benign, with no functional impact. The variants we have studied are intermediate—non-lethal but causing disease of varying severity. Mapping onto the thick filament atomic model reveals a disproportionate number of HCM variants in the small regions of surface contact in the IHM interfaces (BH-FH, BH-S2). Moreover, variants in these locations are also associated with earlier onset and more severe disease, supporting the IHM’s critical role in normal functioning of the heart. There are many fewer pathogenic variants in the tail, almost exclusively in interfaces within the backbone, which tend to cause later onset, less severe HCM. These may cause disease indirectly—by mild disturbance of the delicate interactions that stabilize the IHMs in their off-state.

Our results overall highlight the importance of interfaces in generating a wild-type thick filament structure with normal function. Our findings are consistent with a model in which a direct effect on the IHM has the greatest impact (higher severity with BH/FH and BH/S2 variants), while tail variants indirectly impact the IHM, with less disruption of its structure/function causing milder disease. Other variants are scattered over the heads and may affect activity directly (ATPase site, etc.) or allosterically impact interfaces. These insights, based on the native structure of the human cardiac thick filament, may improve risk stratification of HCM patients, predict the impact of newly discovered pathogenic variants within or excluding interfaces, and aid in the design of drugs for the treatment of cardiomyopathies caused by variants in thick filament proteins.

## Methods

### Sequences

We used the sequences from human cardiac myosin 2, ELC, RLC, cMyBP-C and titin, following ^16^.

### Curation of HCM and DCM variants

We first identified pathogenic (P), likely pathogenic (LP), likely benign (LB) and benign (B) missense variants in MYH7, MYL2, MYL3, MYBPC3 in ClinVar ^36^. We then curated the final list of PLP (P and/or LP; denoted “pathogenic”) and BLB (LB and/or B; denoted “benign”) variants for HCM and DCM by comparing their phenotypes to previously published articles ^2^ (Supplementary Datafile 2). Less than 3% of HCM cases have two pathogenic variants, generally causing worse disease than one mutation ^37 38^. So, usually, only one HCM or DCM variant is present in one person, and if so, because heterozygous, only in one half of MHC, ELC, RLC, cMyBP-C, or titin molecules.

### Mapping of HCM and DCM variants

We used ChimeraX ^39^ to map the curated list of ClinVar variants onto the atomic model of the human cardiac thick filament C-zone (PDB 8G4L).

### Assessing the regional distribution of variants

We determined whether variants were over- represented in a region of interest following our previous approach ^12^: the proportion of variants that fell within an interface region was compared with the proportion that would be expected under a uniform distribution (given by the number of amino acids in the interface / total number of amino acids in the protein or region of interest). Proportions were compared using a binomial test, implemented in R ^12^.

### SHaRe endpoints used for assessment of clinical manifestations of HCM

We selected endpoints reported in (^2^) with updated genotype and longitudinal clinical information. For all data, age indicates mean age ± S.D. for each group, primary diagnosis indicates clinical demonstration of left ventricular wall thickness > 13 mm in adults or z-score based on body mass index for children. Ventricular arrhythmic composite indicates the first occurrence of sudden cardiac death, resuscitated cardiac arrest or appropriate implacable cardioverter-defibrillator therapy. Heart failure composite indicates first occurrences of cardiac transplantation, left ventricular assist device implantation, left ventricular ejection fraction < 35% or New York Heart Association class III/IV symptoms.

## Acknowledgments

Supported by NIH NHLBI HL164560 (RP), NIAMS AR081941 (RP), NHLBI K08HL164885 (YK), the British Heart Foundation’s Big Beat Challenge award to CureHeart (BBC/F/21/220106) (CH, CES, and JGS) the National Science Foundation (NSF) Engineering Research Center on Cellular Metamaterials EEC-1647837 (CES and JGS). See Supplementary Info for details on the SHaRE investigators.

## Authors contributions

Conceptualization: R.C., R.P. Data curation: Y. K., C.E.S., J.G.S., Analysis: D.D., R.P., R.C., Y. K., J.G.S., C.E.S., C.H. Writing: original draft, R.P., editing, RC; final draft, all authors. Funding acquisition: R.C., R.P., J.G.S, C.E.S. Supervision: R.C., R.P.

---

**Supplementary Fig. 1.**
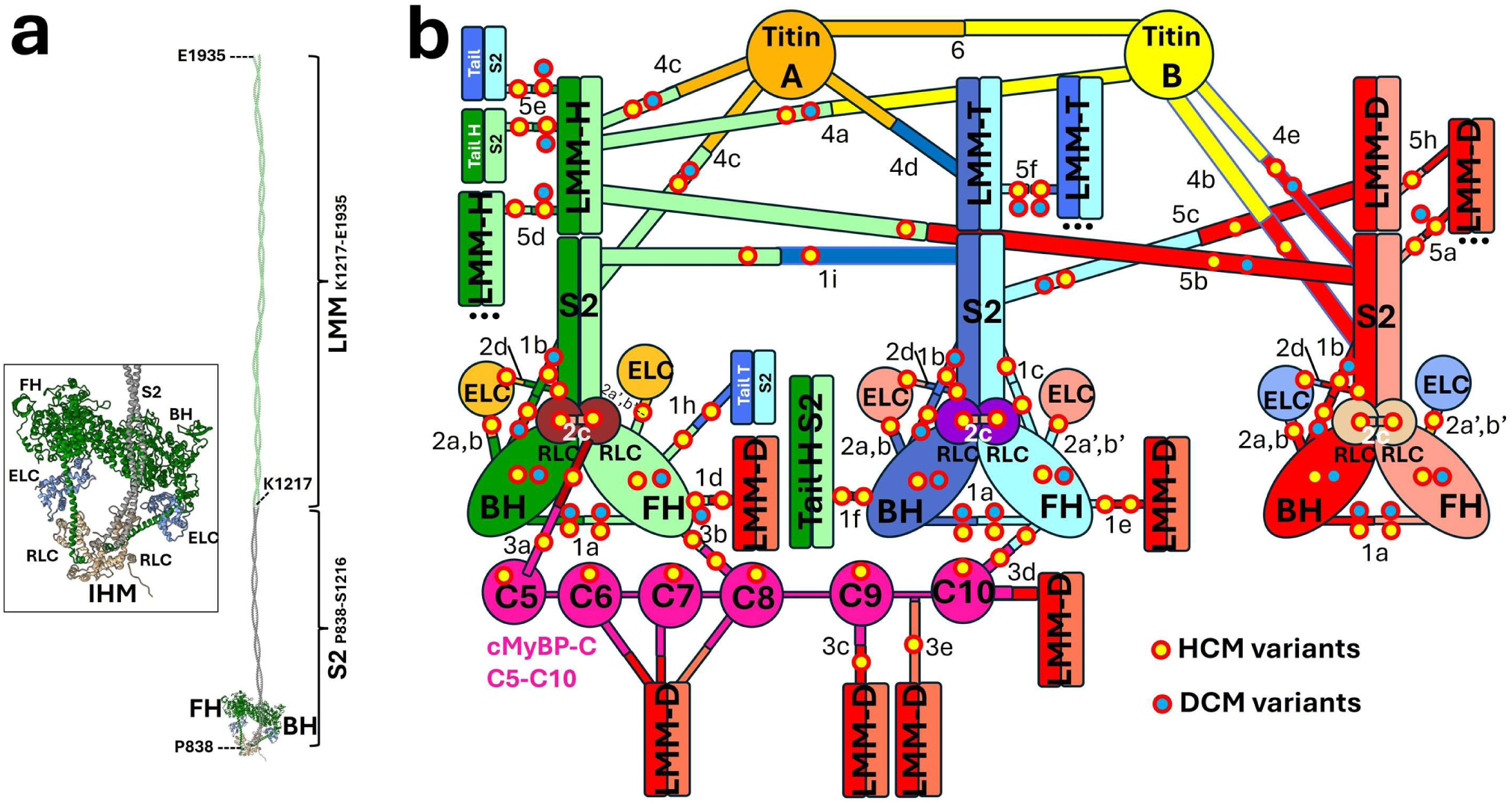
Human cardiac myosin molecule and thick filament interactome. **(a)** The human cardiac myosin molecule is composed of two myosin heavy chains (MHC), whose N-terminal halves form two globular heads and C-terminal halves form a α-helical coiled-coil tail, consisting of subfragment 2 (S2) and light meromyosin (LMM). In the thick filament, the heads interact with each other, creating a blocked head (BH) and free head (FH), and the BH interacts with S2, together forming the interacting-heads motif (IHM). **(b)** Within the thick filament, myosin molecules are assembled with each other, with cMyBP-C and with titin to form a complex interactome, with 32 different types of interfaces. Different classes of interface involve the myosin heads (type 1), light chains (type 2), cMyBP-C (type 3), titin (type 4), and tails (type 5). These are subdivided (1a, 1b, etc.) according to the different interactions within each class. The presence of HCM and DCM pathogenic variants in these interfaces is shown. Benign variants were absent from all interfaces. Color coding as in Fig. 1.

**Supplementary Fig. 2.**
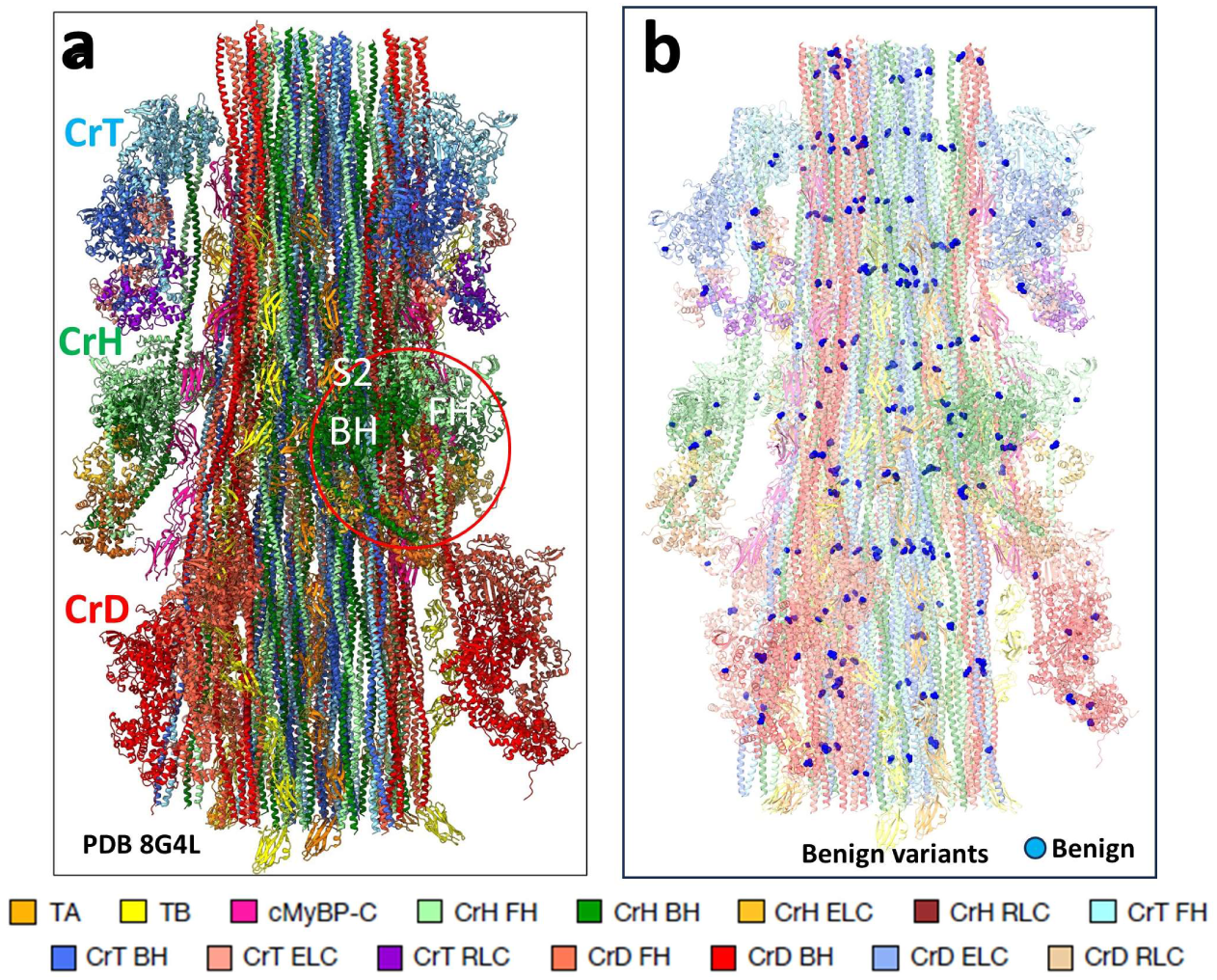
Distribution of benign variants in atomic model. Twenty-one benign variants in the ClinVar curated list are present in the C-zone repeat (PDB 8G4L) **(a, b)**: 11 in MYH7 (two on the MD, one on the lever arm, three on distal S2, five on LMM) and 10 in *MYBPC3* (cMyBP-C domains C6-C9) (Supplementary Tables 2, 3); none are in *MYL3* nor *MYL2*. The benign variants are absent from all filament interfaces. Compare Fig. 1.

**Supplementary Fig. 3.**
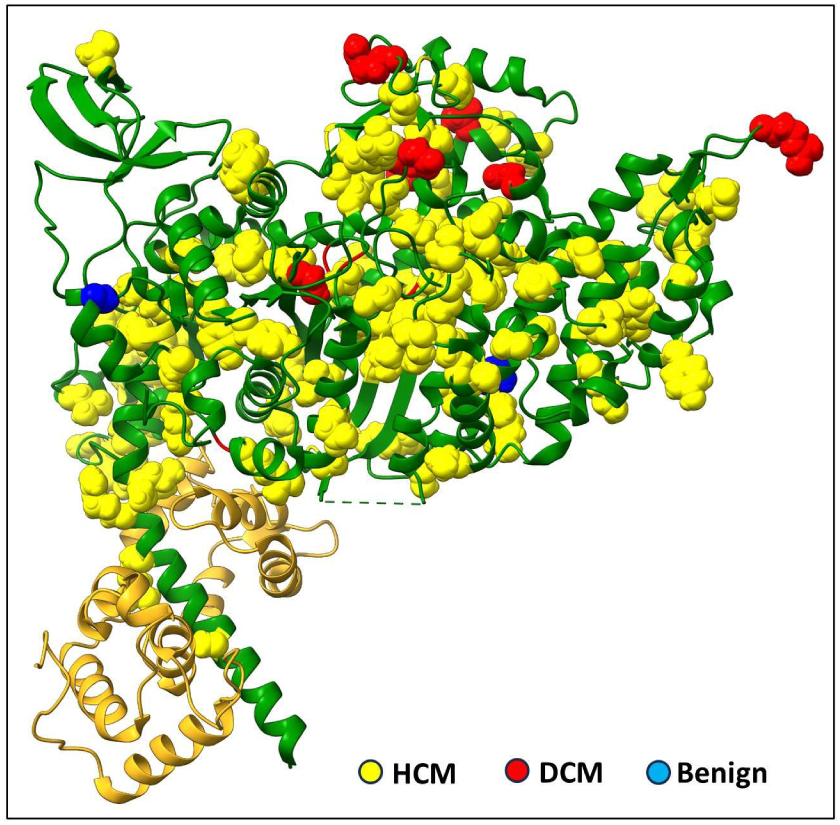
Distribution of curated HCM, DCM and benign variants in the heavy chain of the myosin head. Mapping of the 117 curated HCM variants (yellow), 15 DCM variants (red), and three benign variants (blue) in the CrH BH. CrH FH and CrT, CrD BH/FH would show similar distributions.

**Supplementary Fig. 4.**
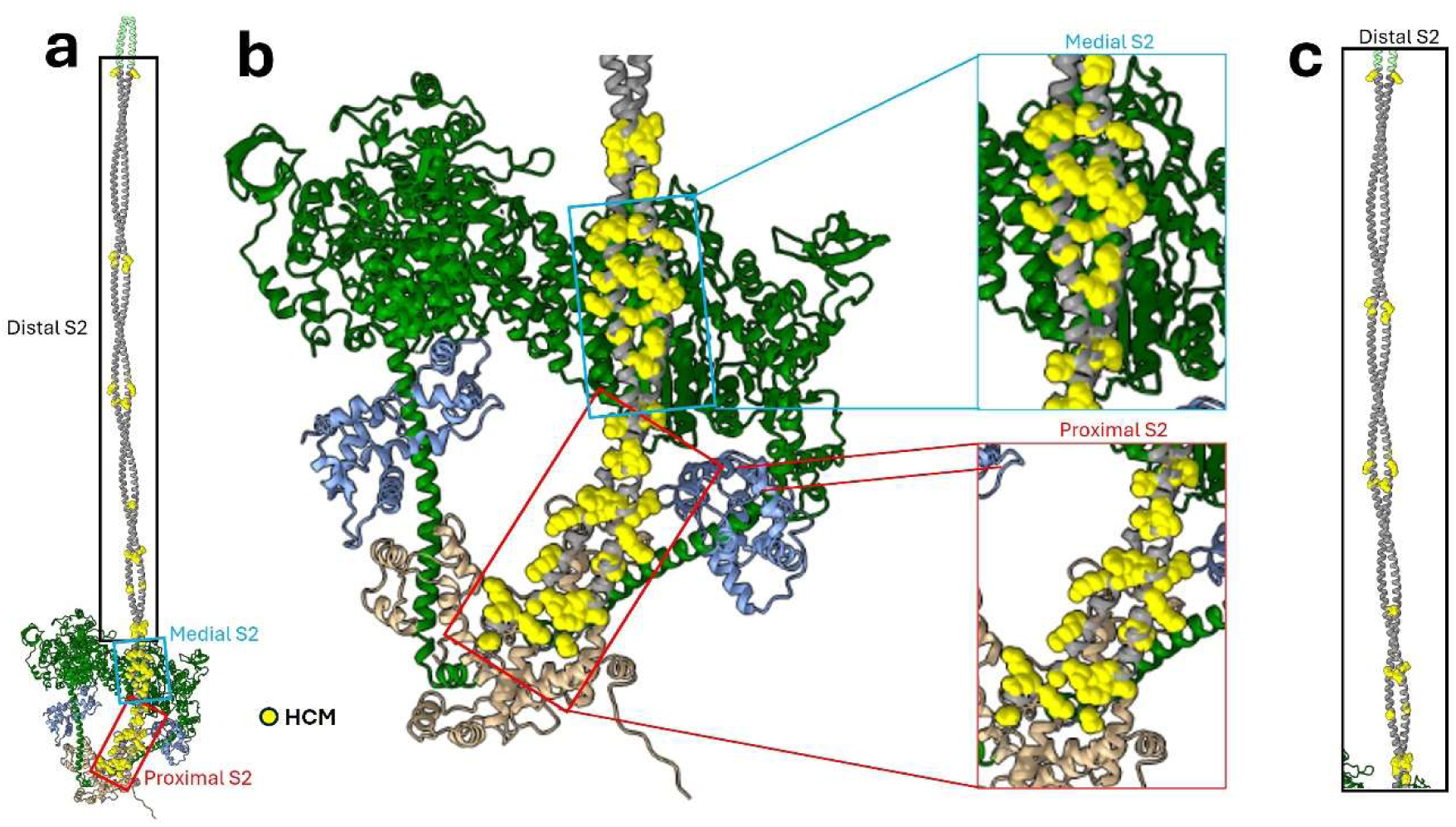
Mapping of HCM variants to the coiled-coil α-helices of S2. (a-c) 37 HCM variants (yellow) (Supplementary Tables 3, 4) are on the proximal (red box), medial (blue box) and distal (black box) regions of CrH S2. Medial S2 is involved in BH-S2 interface type “1b” (Fig. 3, Supplementary Tables 1, 3, 4); proximal S2 is not involved in any interface. Similar distributions occur on tails T and D.

**Supplementary Fig. 5.**
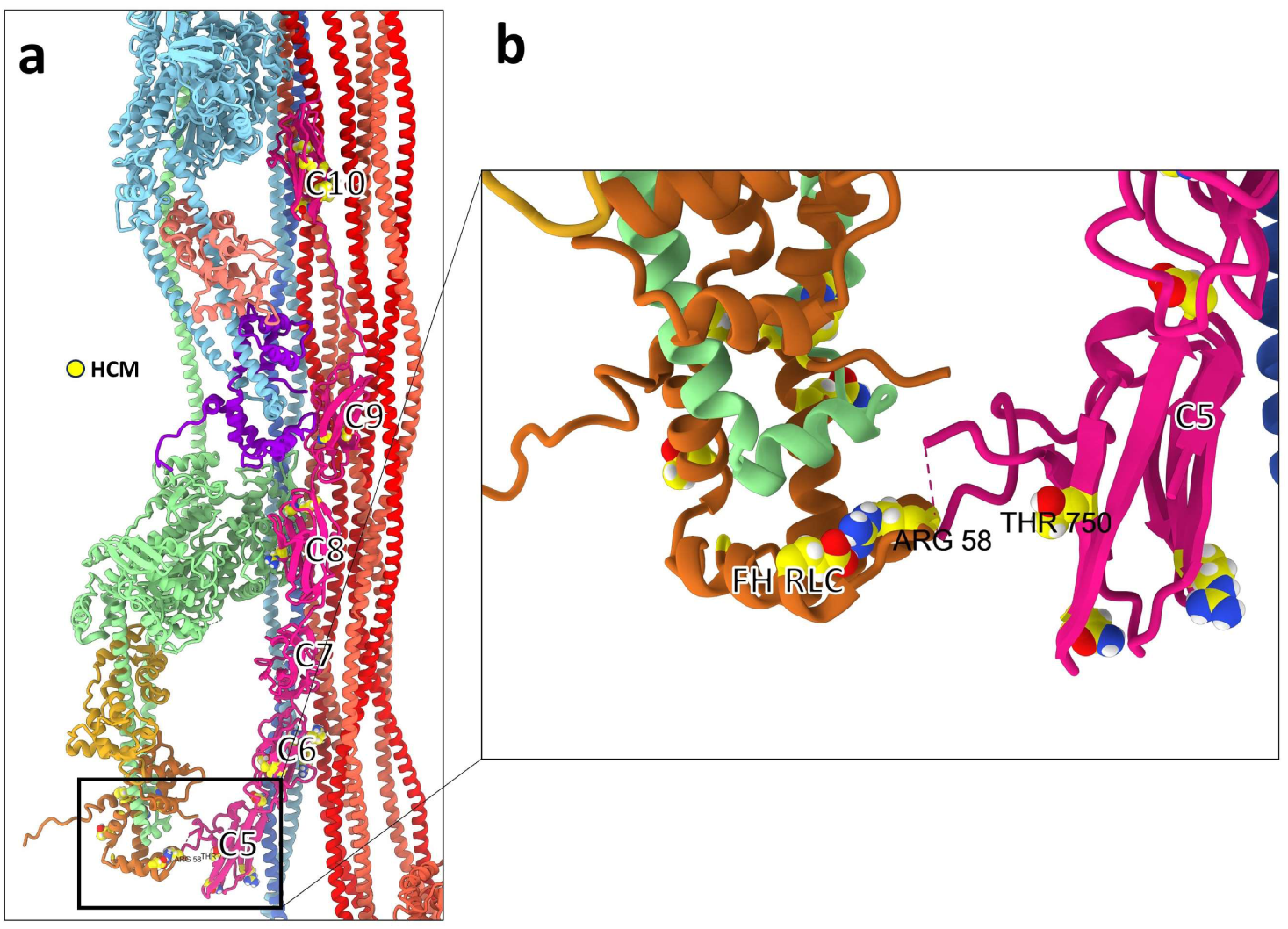
An HCM *MYL2* and a *MYBPC3* domain C5 variant are in the CrH FH – C5 interface. **(a)** Interface type 3a between CrH FH RLC and cMyBP-C domain C5 (black box, zoomed in **b**), showing locations of *MYL2* variant (Arg58Gln) and *MYBPC3* Thr750Met (see red box in Fig. 6a; Supplementary Tables 3, 4).

**Supplementary Fig. 6.**
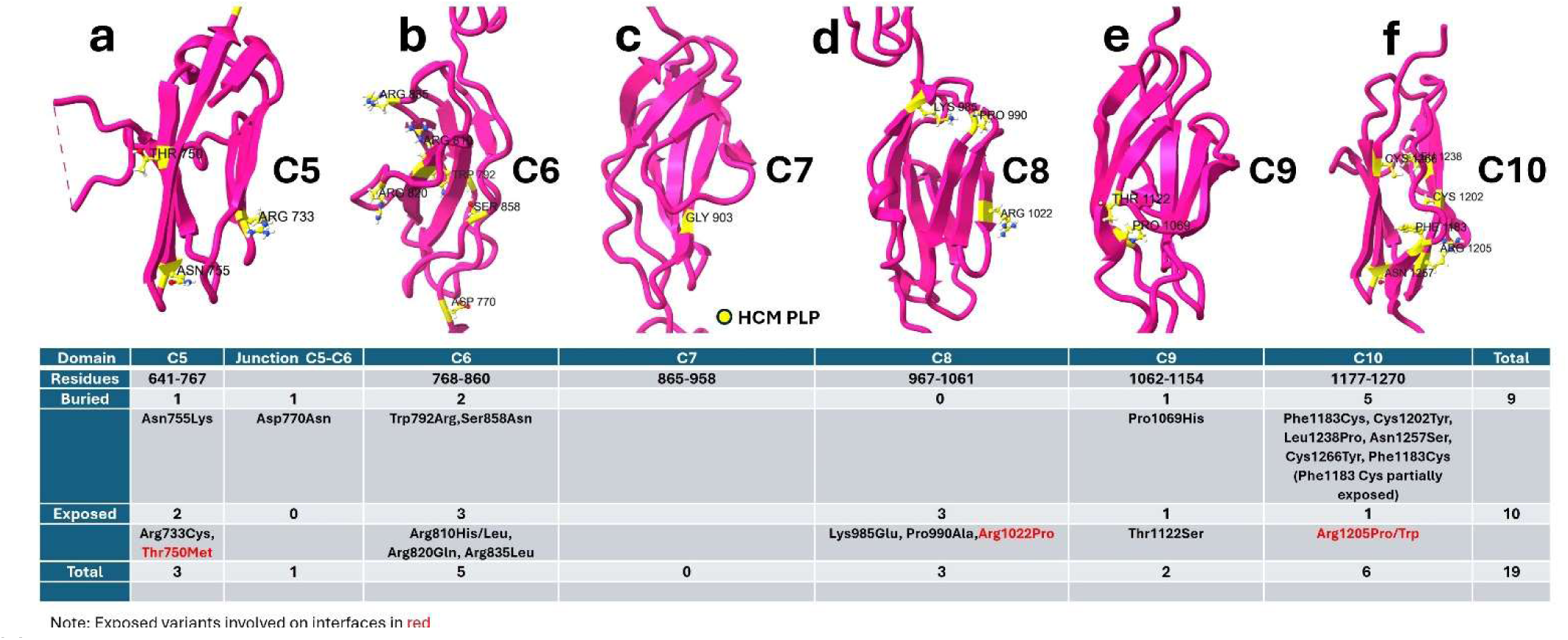
HCM *MYBPC3* variants on cMyBP-C domains C5-C10 are either buried in the domains or exposed on their surface. **(a-f)** Mapping HCM variants on cMyBP-C domains C5-C10 of PDB 8G4L showing buried and exposed variants (see Table). Domains C6 and C10 are hotspots for exposed (C6) and buried (C10) *MYBPC3* variants.

**Supplementary Fig. 7.**
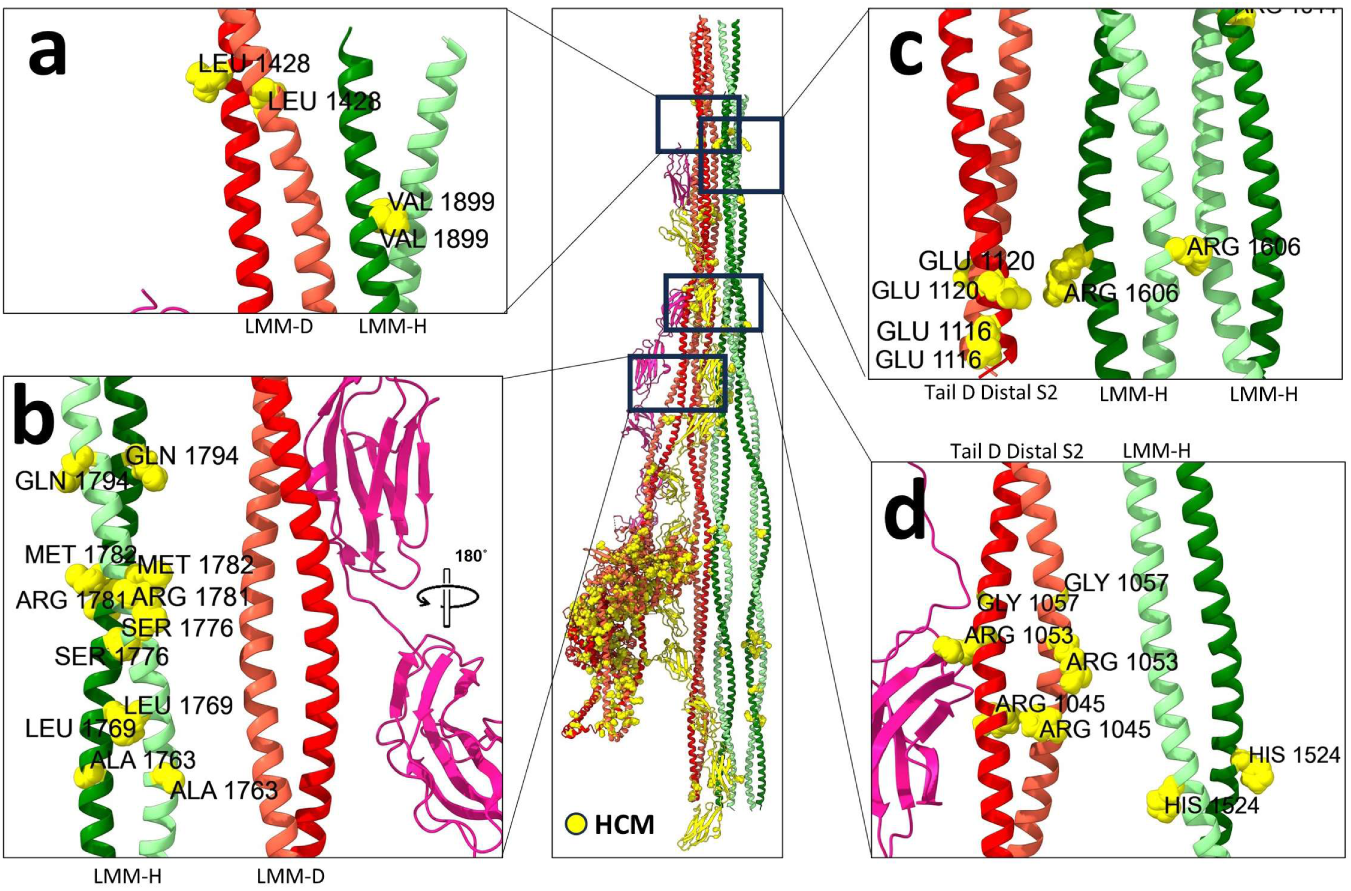
HCM *MYH7* variants are present in LMM – LMM and distal S2 – LMM interfaces. Black boxes in the center image show variant-containing interfaces between (Supplementary Tables 3, 4): **(a)** LMM-D and LMM-H; **(b)** LMM-H and LMM-D, involving a hotspot of six variants on LMM-H with no variants on LMM-D; **(c)** tail D and the same variant (Arg1606Cys) on the middle and right LMM-Hs; **(d)** tail D distal S2 and LMM-H. Some of these variants, which are charge-changing or helix-breaking, could impair interfaces holding LMM-Ds onto the LMM-H backbone shell.

**Supplementary Fig. 8.**
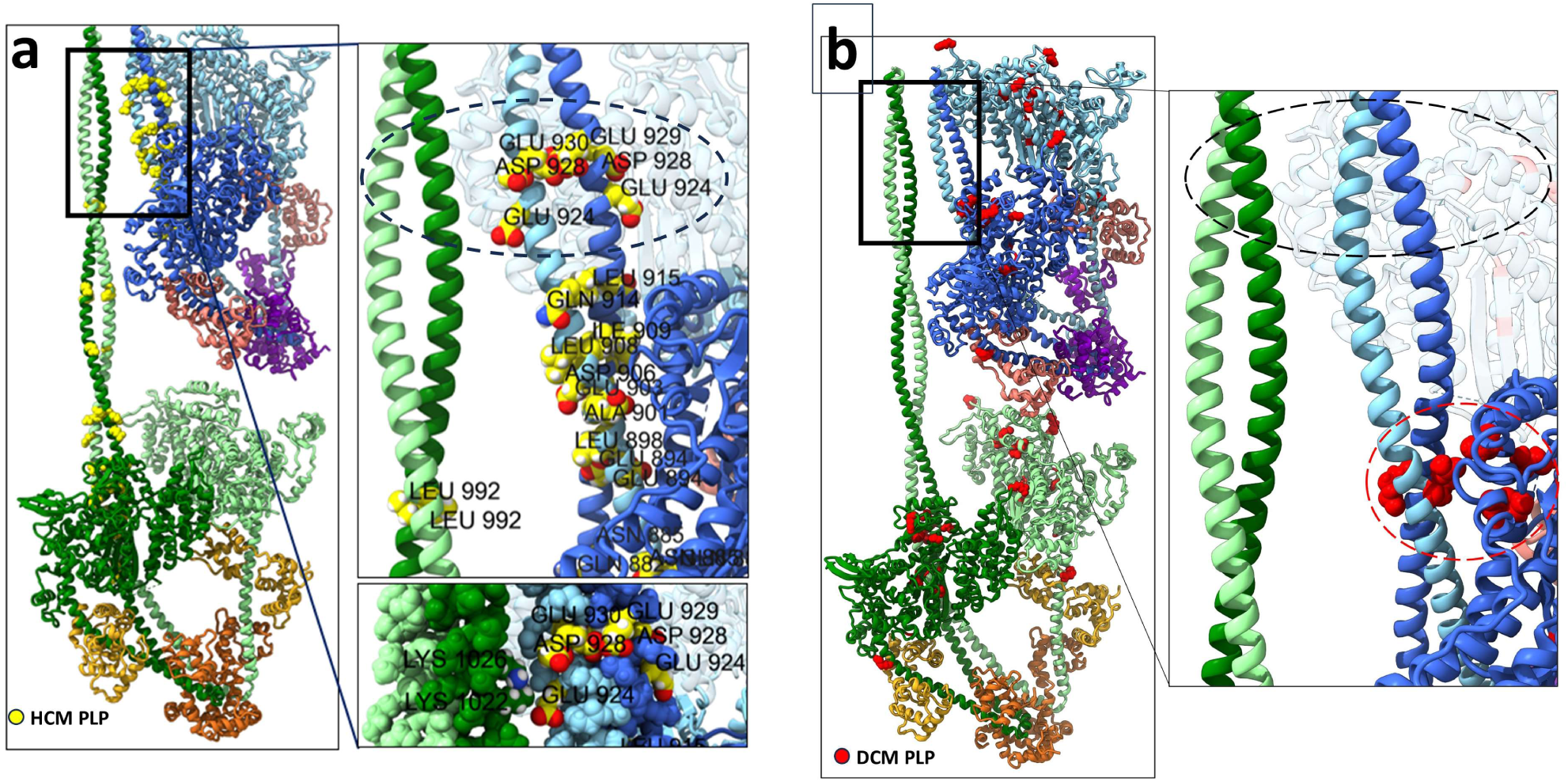
HCM -but not DCM- variants are on interfaces between the distal S2s of tails H and T. **(a)** black box shows HCM variants (yellow) on tail T distal S2 (blue; Ring 2 residues 920-935) at its interface with tail H distal S2 (green), zoomed on right (ellipse, type “5g”, Supplementary Tables 1, 3, 4), showing the two contacts involving HCM variants at residues Glu924Lys and Asp928Asn/Val (bottom inset). **(b)** shows, in contrast, that in the same region (ellipse) as in **(a)**, there are no DCM variants (red).

**Supplementary Fig. 9.**
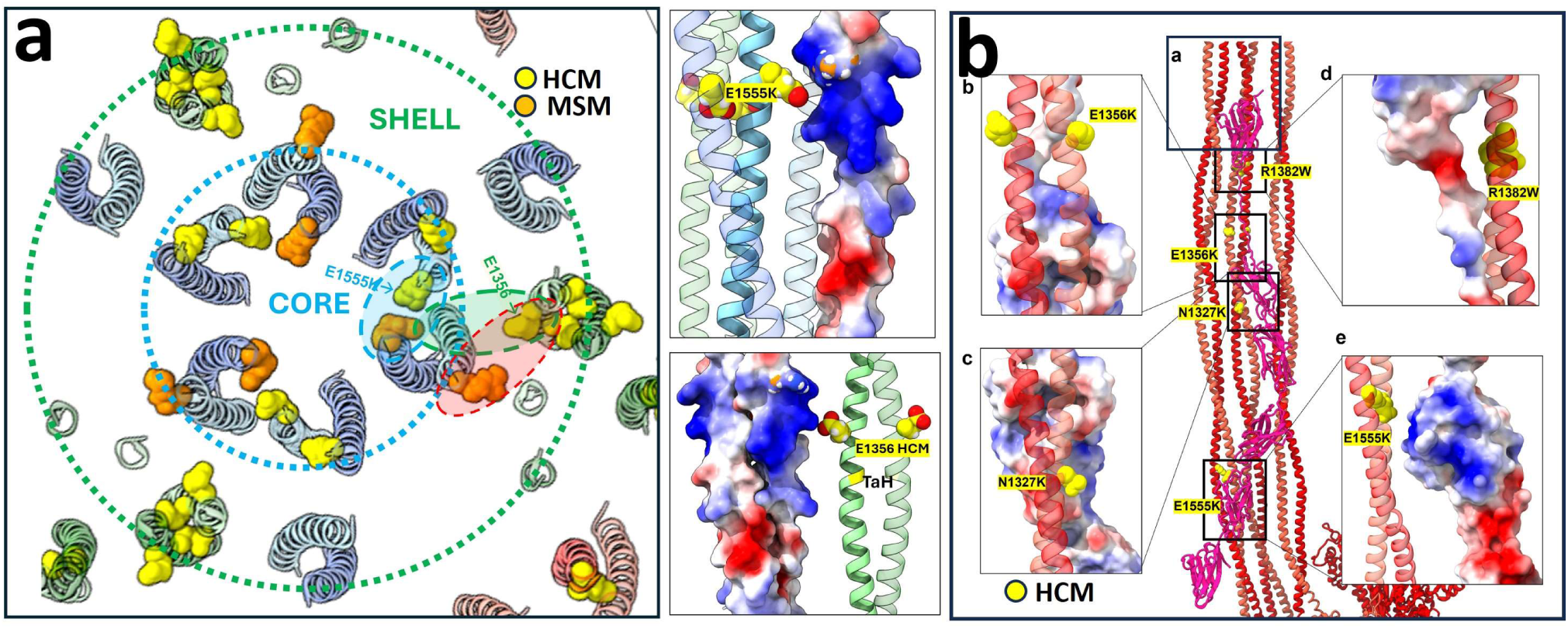
Mapping of HCM variants on LMM-LMM interfaces. **(a)** HCM variant Glu1356Lys (yellow spheres) on LMM-H (green α-helices) will impact LMM-H – LMM-T interface type 5e (Supplementary Tables 1, 3, 4; green ellipse) involved in backbone shell (green dotted circle) assembly. Glu1555Lys (yellow spheres) on LMM-T (blue α-helices) will impact LMM-T – LMM-T interface type 5f (Supplementary Tables 1, 3, 4; blue ellipse) involved in backbone core (blue dotted circle) assembly. The change in these variants from negative (shown) to positive will convert attractive to repulsive electrostatic interactions (insets at right; blue positive, red negative). These variants reduce myosin incorporation in muscle ^40^, supporting our suggestion of their impact on filament assembly. Interestingly, charge-changing variant Arg1845Trp (orange spheres) in tail T (blue α-helices), could affect <u>simultaneously</u> the formation of the backbone core (blue ellipse) and backbone shell (red ellipse). This variant leads to myosin storage myopathy (MSM) in skeletal muscle ^28^. **(b)** Charge-changing or charge-reversal variants Glu1356Lys, Arg1382Trp, Asn1327Lys and Glu1555Lys on LMM-Ds (red α-helices) could impair electrostatic interactions between these tails (type 5h, Supplementary Table 1, boxes and zoomed insets), which together form a docking platform for cMyBP-C domains C6-C10 (pink) on the filament surface ^16^. Interference with this docking region could compromise cMyBP-C’s proposed role in length-dependent activation (LDA, ^16^. Thus, Glu1356Lys on LMM-H and Glu1555Lys on LMM-T could impair backbone shell and core assembly **(a)**, while in LMM-D the same variants could, in addition, impair LDA.

**Supplementary Fig. 10.**
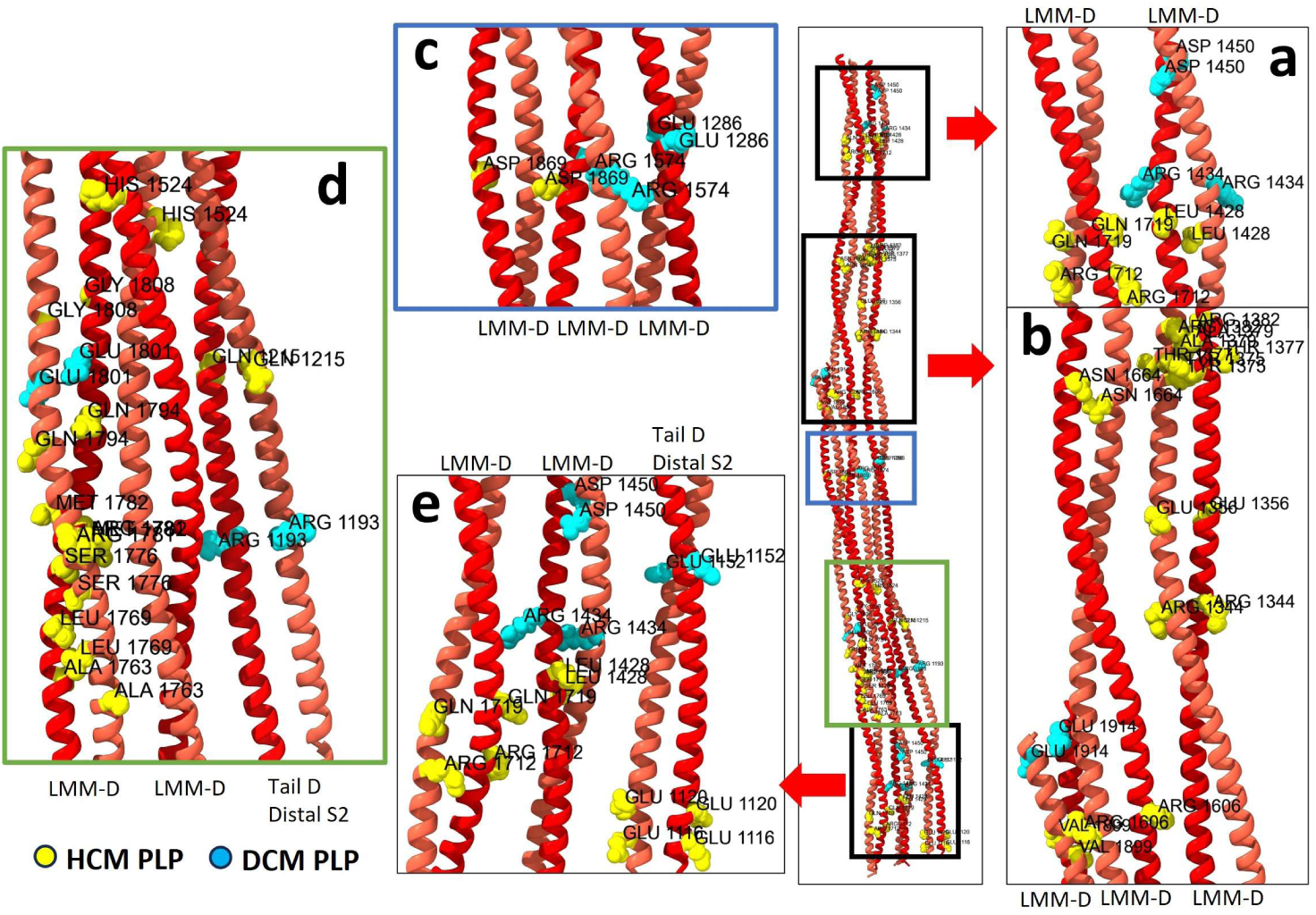
HCM and DCM variants on tail-D could destabilize the tail-D sheets involved in cMyBP-C binding. Five groups of HCM and DCM variants (yellow and blue spheres, respectively) are shown in colored boxes and zoomed for detail. The variants (Supplementary Tables 3, 4) are in pairs (one copy on each α-helix of a tail), that could destabilize the seam of the coiled-coil α-helices, and/or affect the inter-tail-D interactions and thus tail-D sheet stability, key to cMyBP-C binding.

**Supplementary Fig. 11.**
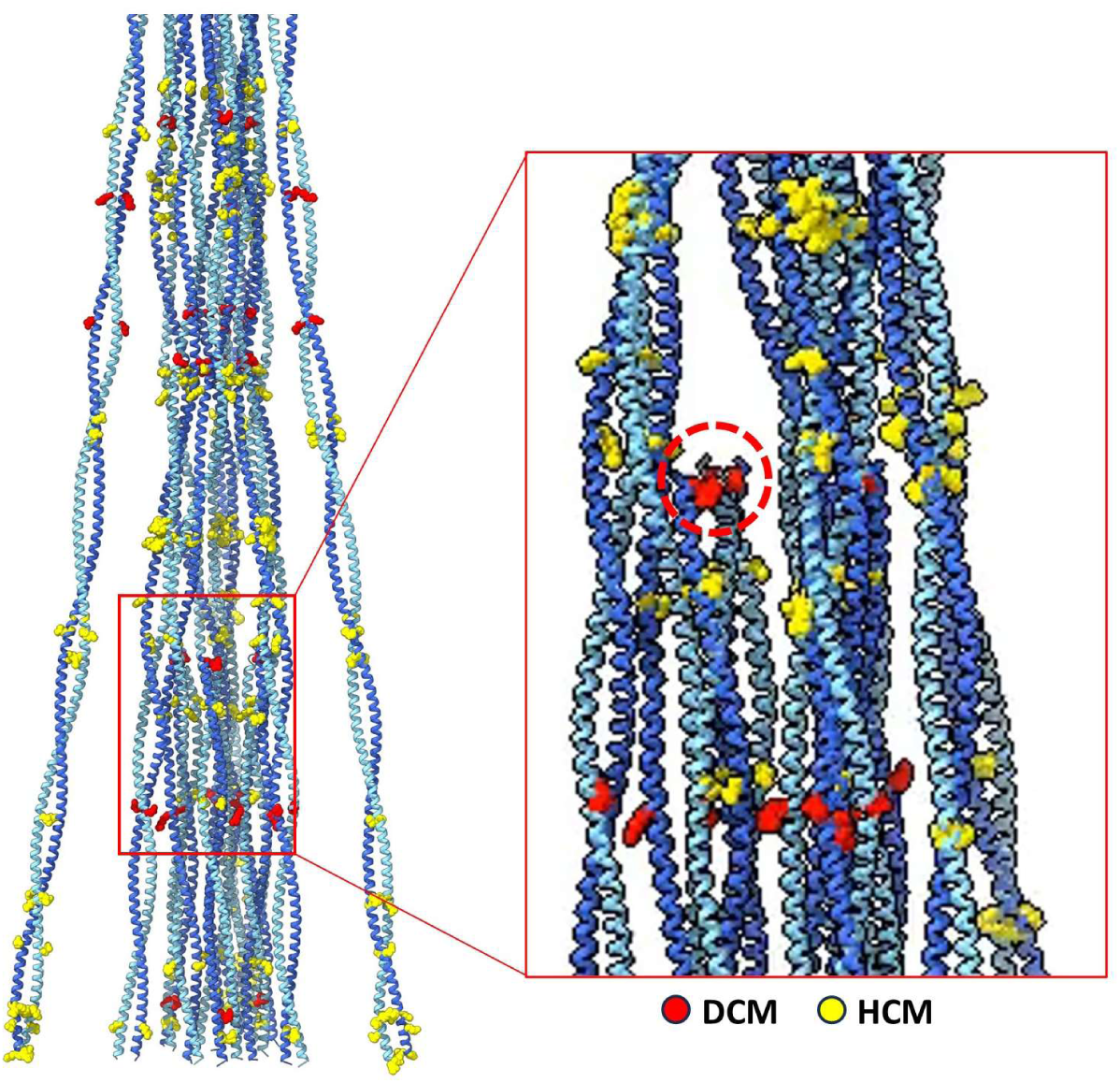
Mapping HCM and DCM variants on tail T interfaces. **Left:** DCM and HCM variants (red and yellow, respectively) are mapped onto tail T in the backbone core. Because 3 identical tails, each with identical variants in the 2 α-helical chains, come close to each other in axial register in the core of the 3-fold symmetric filament ^16^, each variant creates a horizontal stripe of 6 identical residues, which repeats at 430 Å intervals. HCM and DCM variants are on different “stripes” due to their different locations along the tail. **Right:** Red box is slightly rotated and zoomed to show DCM variant at LMM-T C-terminal (red circle).

**Supplementary Fig. 12.**
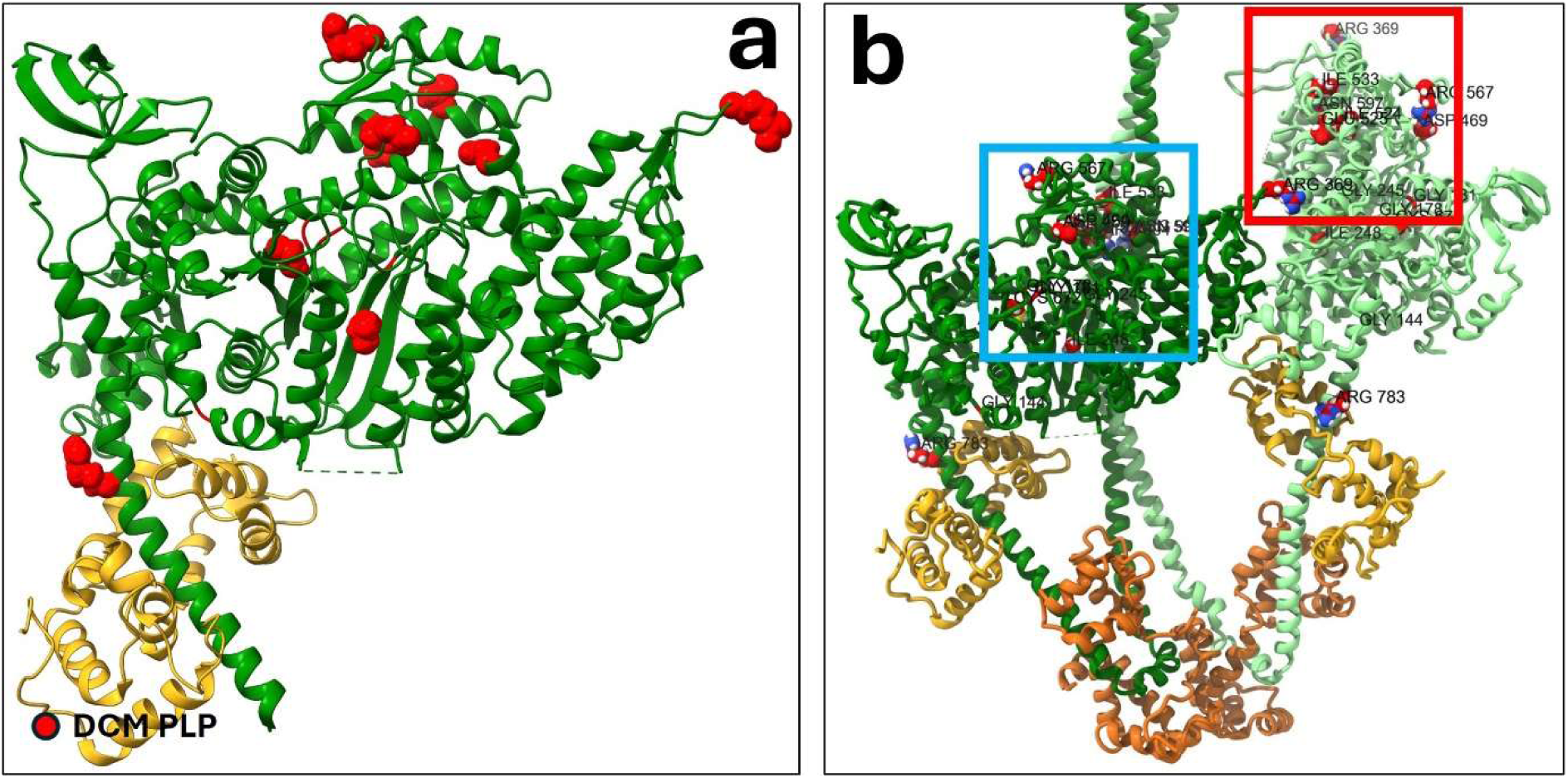
Fifteen DCM *MYH7* variants are on the myosin head, three in IHM interfaces. **(a)** Some of the 15 DCM curated variants are shown on the CrH BH (red spheres). A similar distribution occurs on the CrH FH, and the BH and FH of CrT and CrD. **(b)** The three variants on interfaces are: Arg369Gln in both the BH-FH (type 1a, red box) and FH - LMM-D interfaces (type 1d, Supplementary Table 4); Glu525Lys (together with DCM Arg904Pro on tail H medial S2) in the BH-S2 interface (type 1b, blue box); and Arg783Pro in the BH and FH lever arm – ELC interface (type 2b). Interestingly Arg783 in the same interface but differently mutated (Arg783Cys/His) leads to HCM. The remaining 12 DCM variants in the head are not in interfaces.

**Supplementary Fig. 13.**
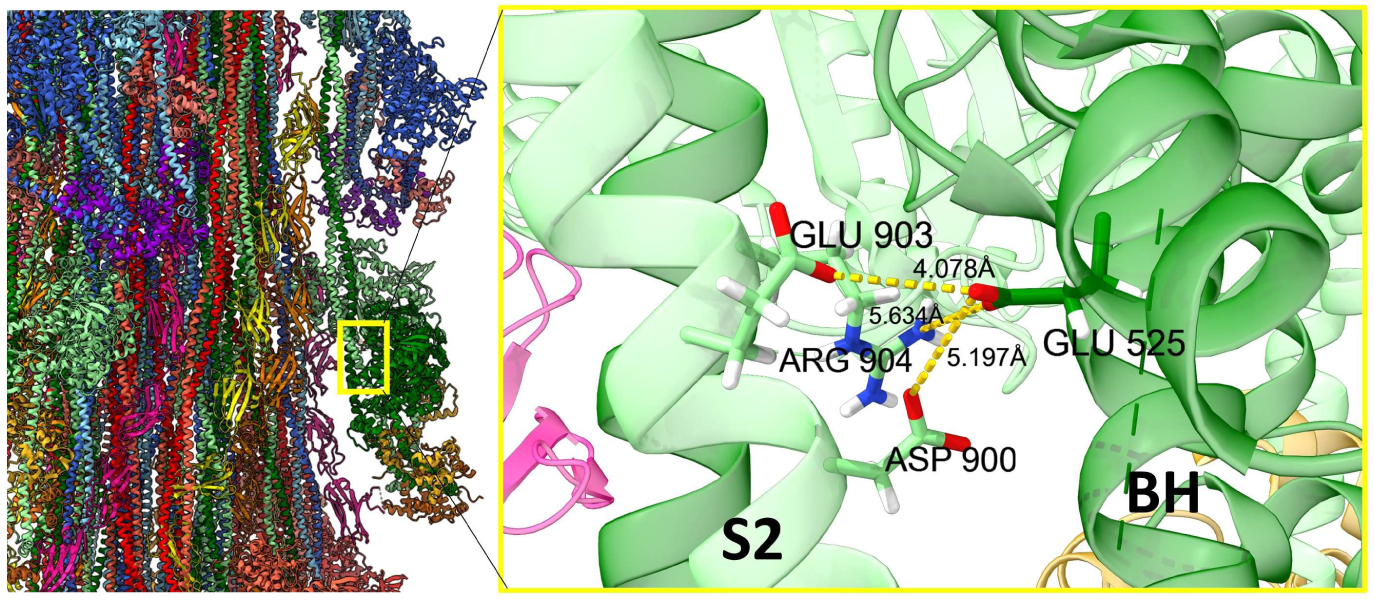
Two HCM and two DCM variants in *MYH7* create an HCM/DCM hotspot on the CrH BH-S2 interface. Of the 25 DCM pathogenic variants, only two (Glu525Lys on BH MD, Arg904Pro in S2) are in the BH-S2 interface (zoomed from atomic model 8G4L on left). Interestingly, Glu903Gly and Asp906Gly are charge-decreasing variants that cause HCM. This HCM-DCM hotspot suggests different – but closely-related – structural origins for these two dissimilar diseases caused by variants in the same interaction interface. Higher resolution studies could open the way to understanding the structural origin of this dissimilarity. A similar hotspot occurs on the CrT IHM.

**Supplementary Fig. 14.**
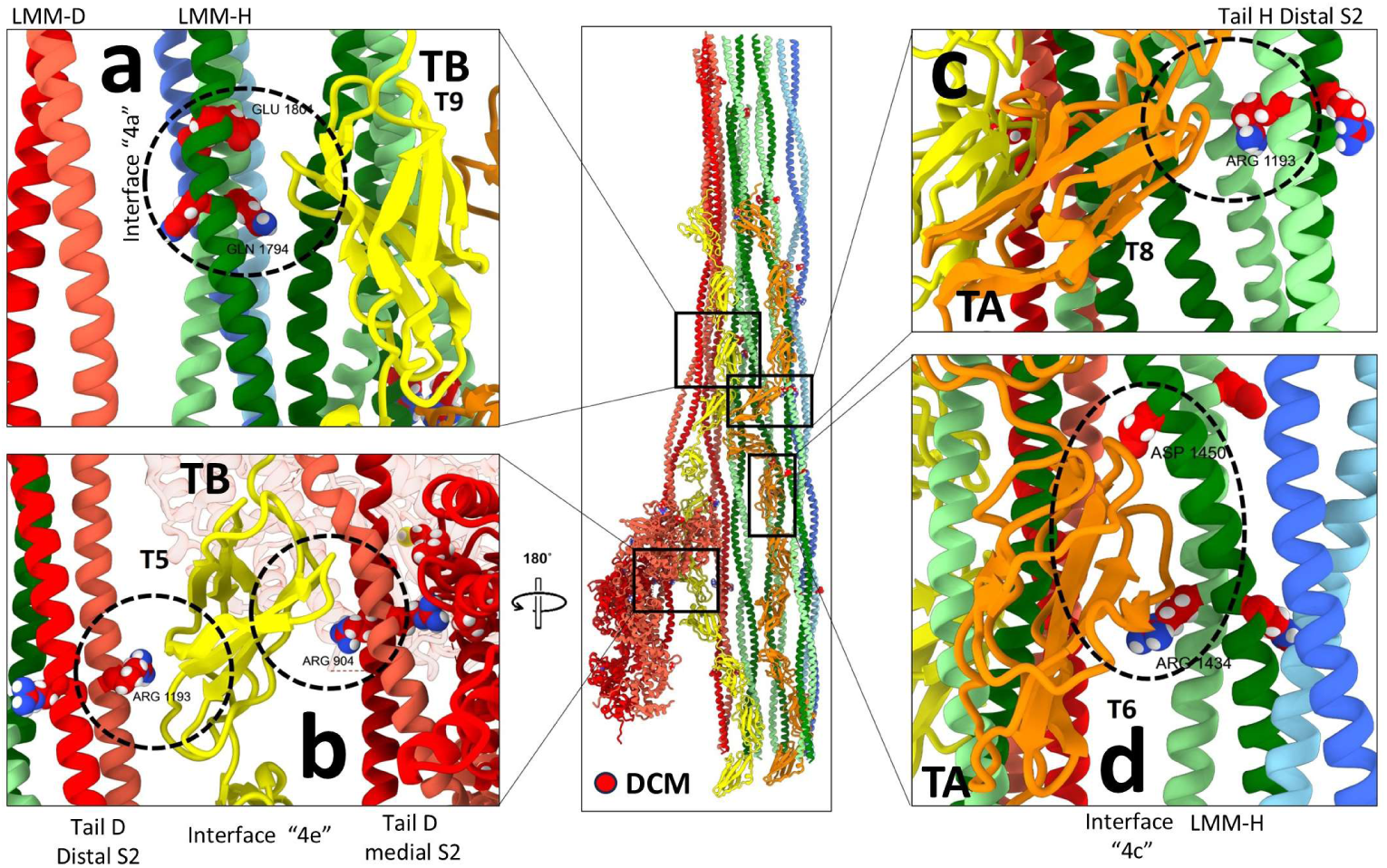
DCM *MYH7* variants occur in titin-S2 and titin-LMM interfaces. **(a)** LMM-H – TB interface (type 4a): Glu1801Lys and Gln1794Glu DCM variants impair the TaH – TB(T9) interface. **(b)** Tail D S2 – TB interface (types 4b, 4e): Arg1193His and Arg904Cys/His DCM variants impair tail D S2 – TB(T5) interface. Note DCM Arg904Cys/His is also involved in HCM-DCM hotspot on BH-S2 interface “1b” (Supplementary Fig. 13). **(c)** Tail H distal S2 – TA interface (type 4c): the Arg1193His DCM variant impacts the tail H distal S2 – TA(T8) interface. **(d)** LMM-H – TA interface (type 4c): Arg1434Cys and Asp1450Asn DCM variants impair the LMM-H – TA(T6) interface.

**Supplementary Fig. 15.**
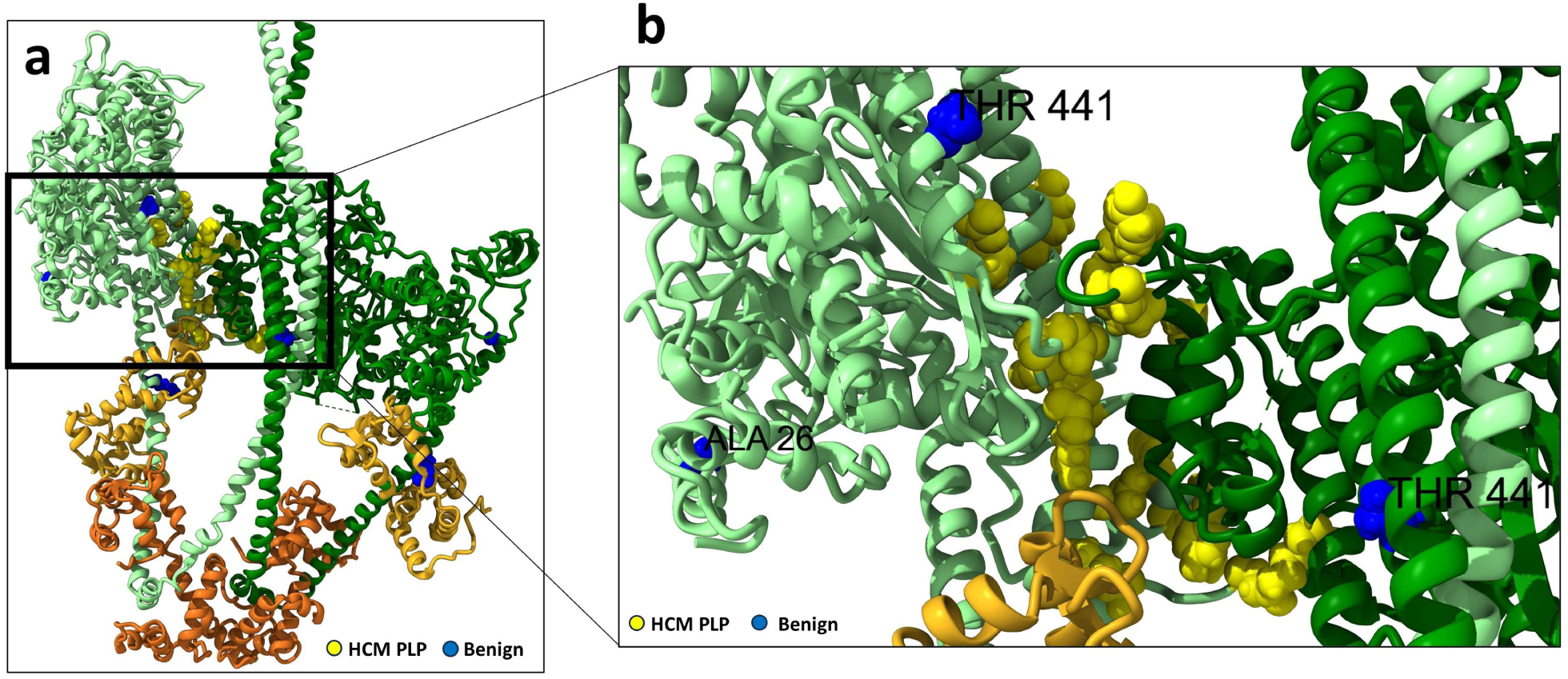
Benign variants on myosin head are absent from interfaces. **(a)** Of the three benign variants in the myosin head (blue), two are in the MD (Ala26Val, Thr441Met) and one on the lever arm (Arg787His). None are in the BH-FH interface, where 24 HCM variants (yellow) are present, zoomed in **(b)**. Shown here for CrH: similar for CrT.

**Supplementary Fig. 16.**
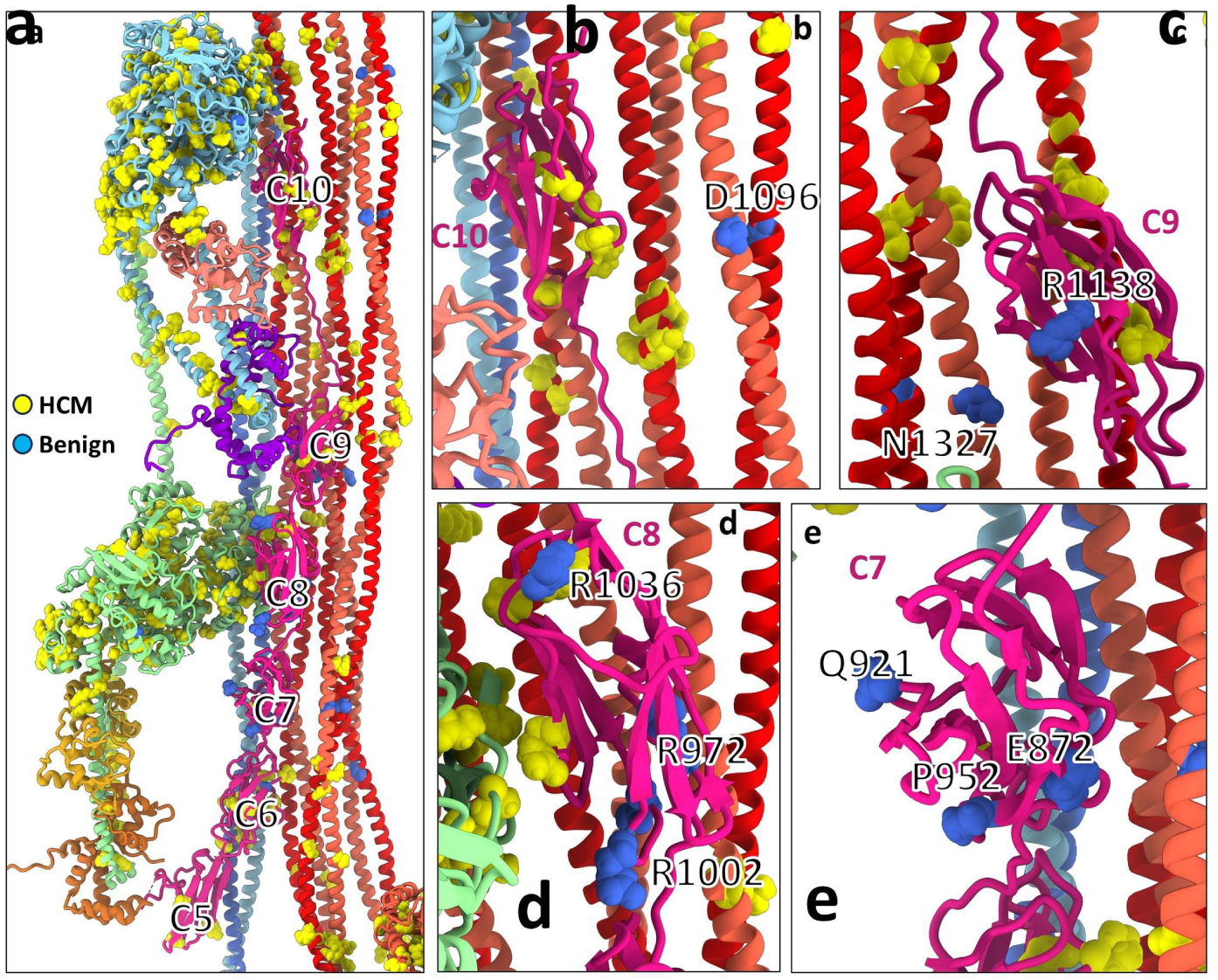
Benign *MYH7* and *MYBPC3* variants are absent from interfaces in the C-zone. **(a)** Benign (blue) and HCM (yellow) variants on cMyBP-C domains and on tail D S2 and LMM-Ds. **(b)** Benign *MYH7* variant Asp1096Tyr on tail D distal S2 is not involved in tail D – tail D interactions while HCM variants are. **(c)** Benign *MYBPC*3 variant Arg1138His on C9 is not involved in any interface. **(d)** Three benign *MYBPC3* variants on surface of C8 (Arg1036Cys, Arg972Trp, Arg1002Trp) are not involved in C8 – CrH FH interface, while MYBPC3 HCM variant (Arg1022Pro) is. **(e)** Three benign MYBPC3 variants (Glu872Lys, Gln921Glu, Pro952Ala) in C7 are not involved in any interface.

**Supplementary Fig. 17.**
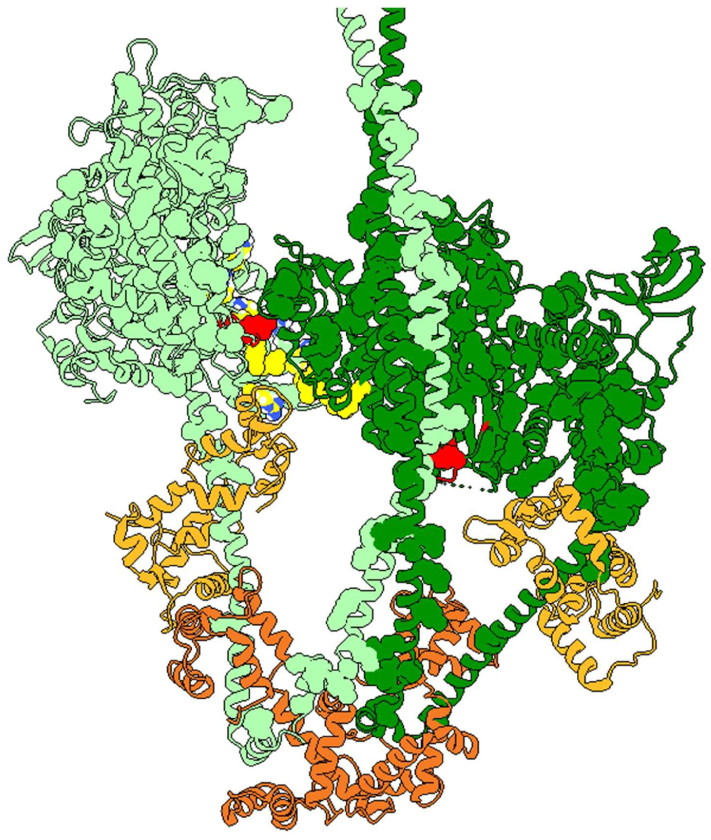
Two *MYH7* HCM variants are simultaneously in the BH-FH and BH-S2 interfaces. Of the 24 variants in the BH-FH interface (yellow), two (red) – Arg249Gln, Arg453Cys/His/Ser are also on the BH-S2 interface (Fig. 3), suggesting a dual IHM destabilizing effect. Shown here for CrH: similar for CrT.

**Supplementary Fig. 18.**
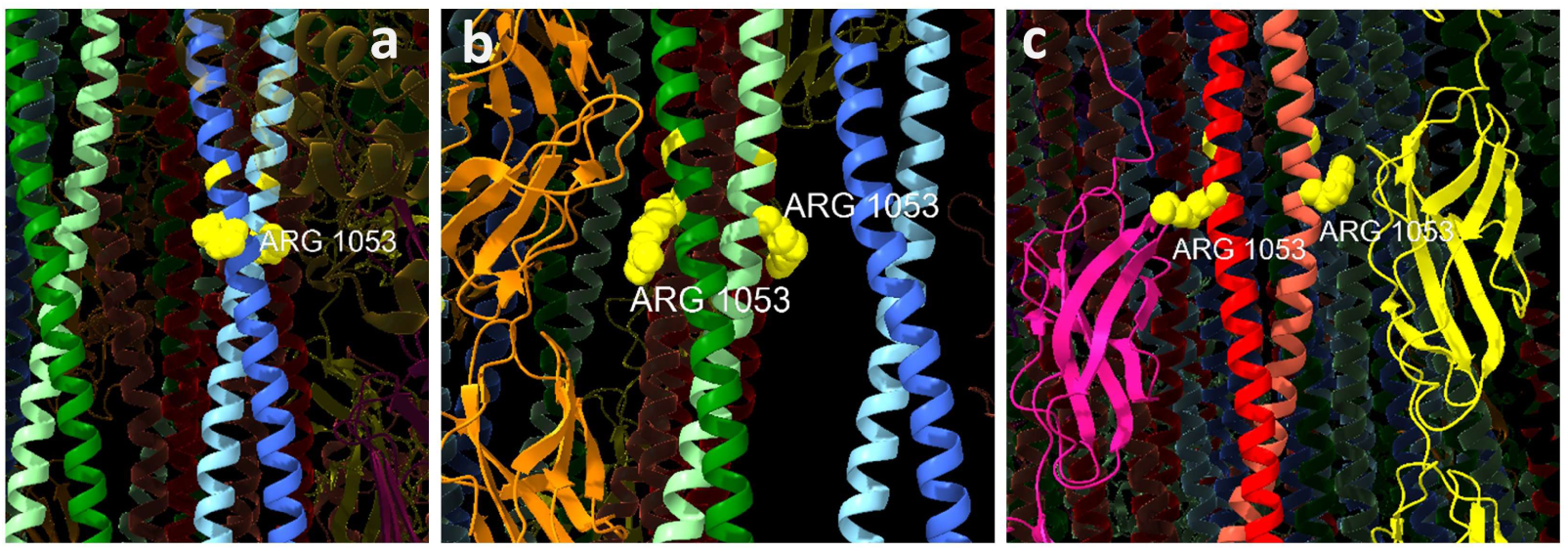
Example of the same HCM mutation in multiple different interfaces. **(a)** Arg1053Gln in LMM-T (blue) in possible interaction with LMM-H (green). **(b)** In LMM-H, Arg1053 interacts with TA (orange) and LMM-T. **(c)** In LMM-D (red), Arg1053 interacts with both cMyBP-C C9 (pink) and TB (yellow). In all cases, Arg1053Gln affects the shell and surface but not the core of the filament.

**Supplementary Fig. 19.**
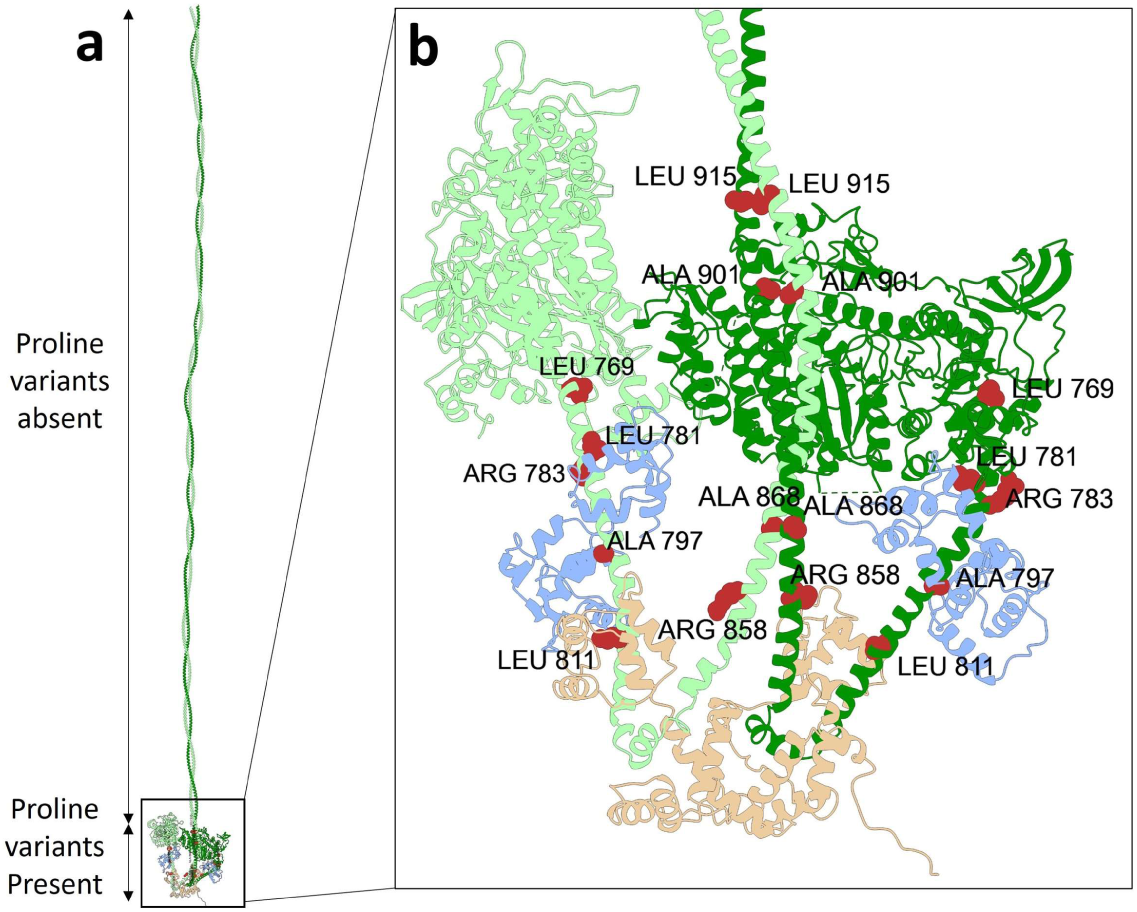
Residues mutated to proline occur only in heads and proximal and medial S2 of IHM, not in distal S2 or LMM. Proximal and medial S2 are part of the IHM, which is not significantly involved in filament assembly but is involved in ATPase activity and the SRX state. Filaments with proline variants in the IHM (red) could therefore still be formed but may not function properly due to impairment of SRX or myosin ATPase. In the main part of the tail (distal S2 and LMM), involved in filament assembly, mutation to proline would cause disruption of α-helices, and interfere with proper filament formation; in this case, proline mutation is likely to be lethal, explaining its absence from ClinVar.

**Supplementary Table 1.**
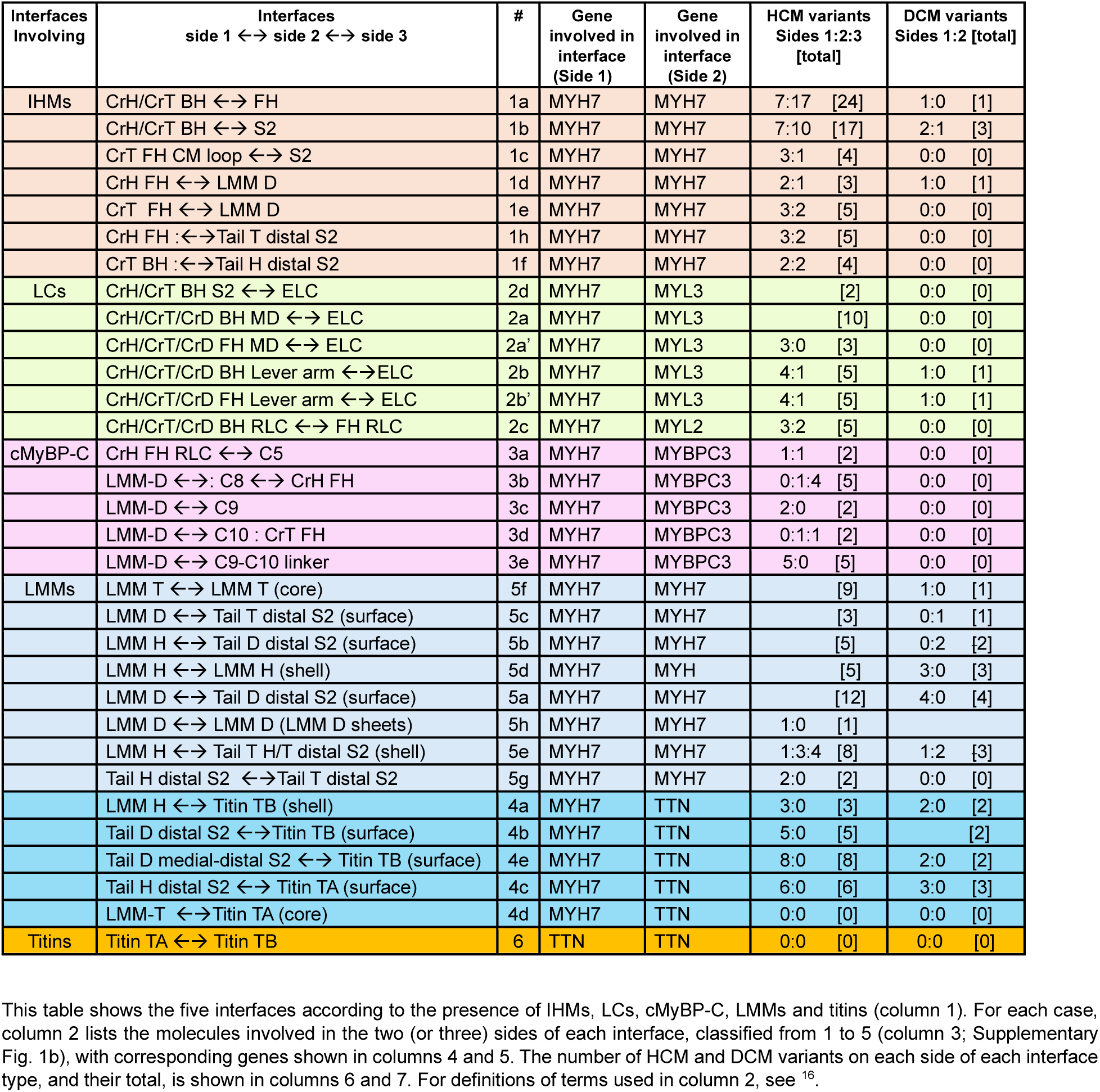
HCM and DCM variants on the 32 interfaces on the human cardiac thick filament.

**Supplementary Table 2.**
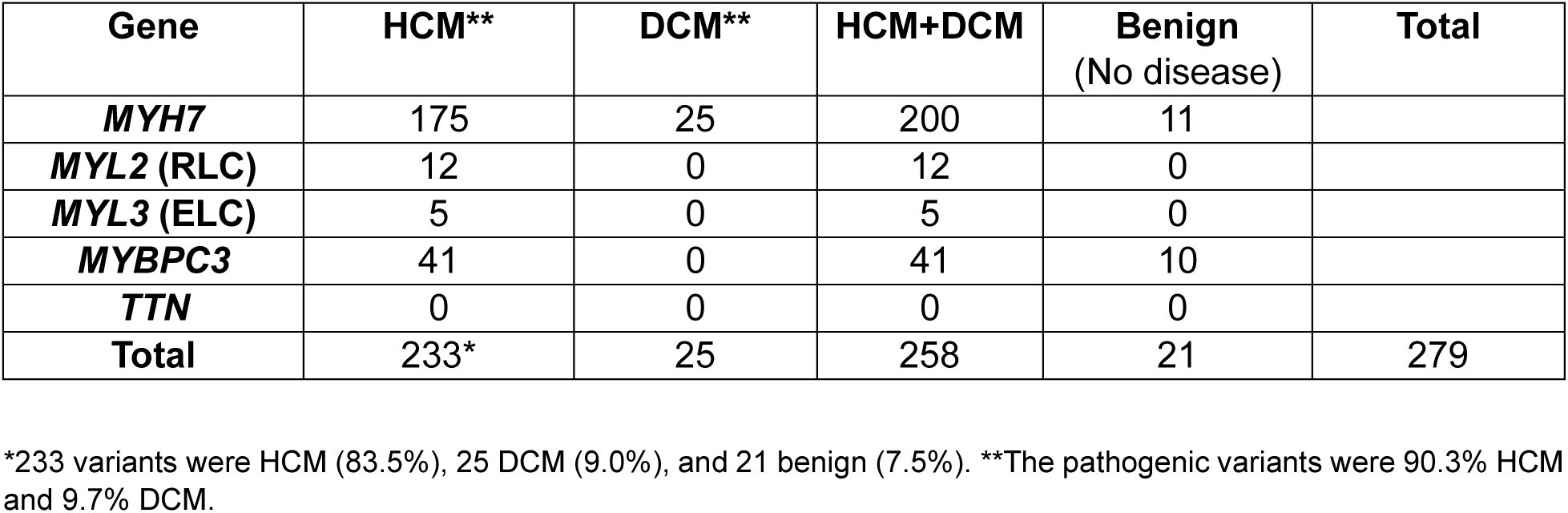
HCM, DCM, and benign *MYH7*, *MYL2/3*, *MYBPC3*, and *TTN* variants curated from ClinVar.

| **Gene** | **HCM\*\*** | **DCM\*\*** | **HCM+DCM** | **Benign**<br>**(No disease)** | **Total** |
| --- | --- | --- | --- | --- | --- |
| ***MYH7*** | 175 | 25 | 200 | 11 |  |
| ***MYL2* (RLC)** | 12 | 0 | 12 | 0 |  |
| ***MYL3* (ELC)** | 5 | 0 | 5 | 0 |  |
| ***MYBPC3*** | 41 | 0 | 41 | 10 |  |
| ***TTN*** | 0 | 0 | 0 | 0 |  |
| **Total** | 233* | 25 | 258 | 21 | 279 |

\*233 variants were HCM (83.5%), 25 DCM (9.0%), and 21 benign (7.5%). \*\*The pathogenic variants were 90.3% HCM and 9.7% DCM.

**Supplementary Table 3.**
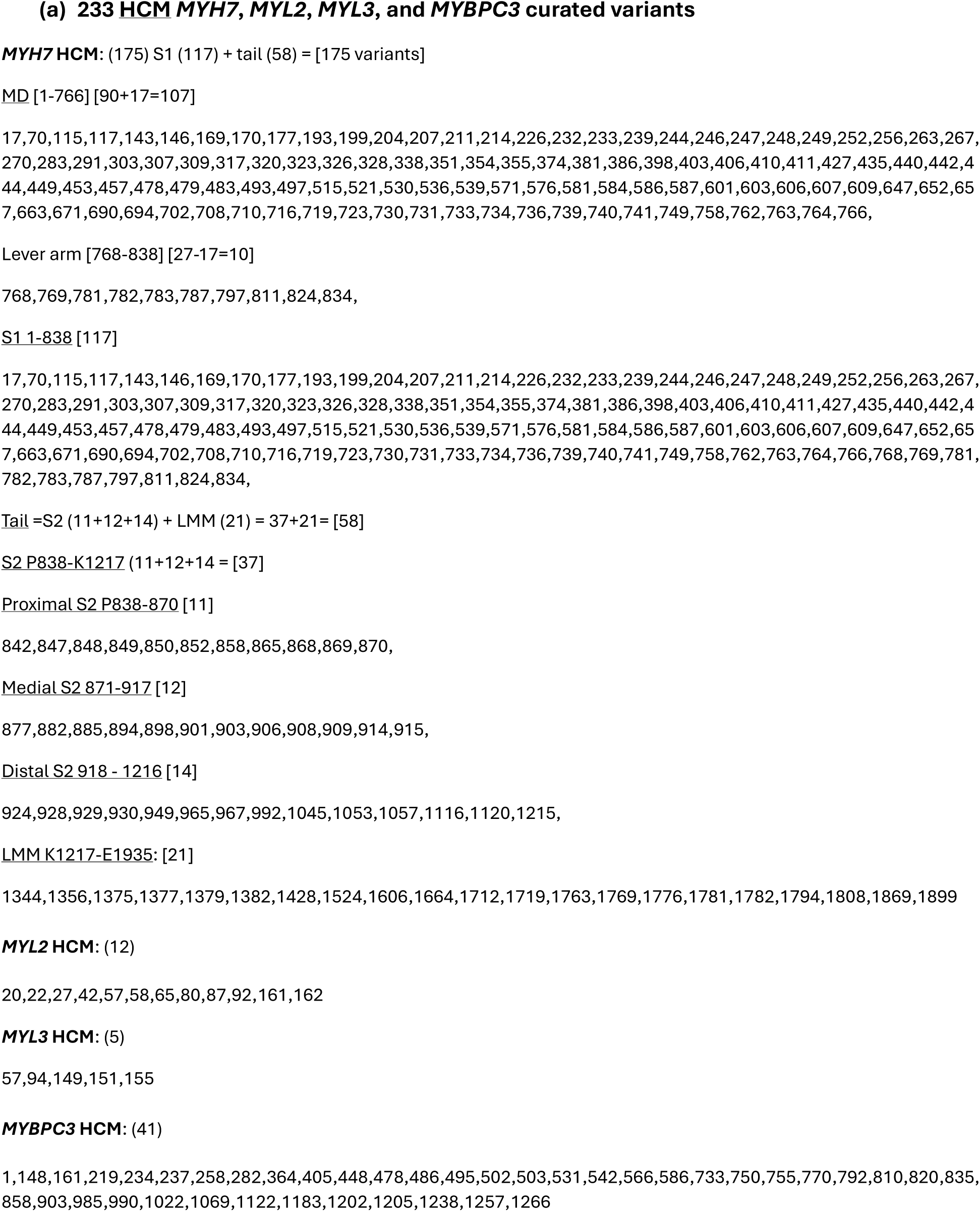
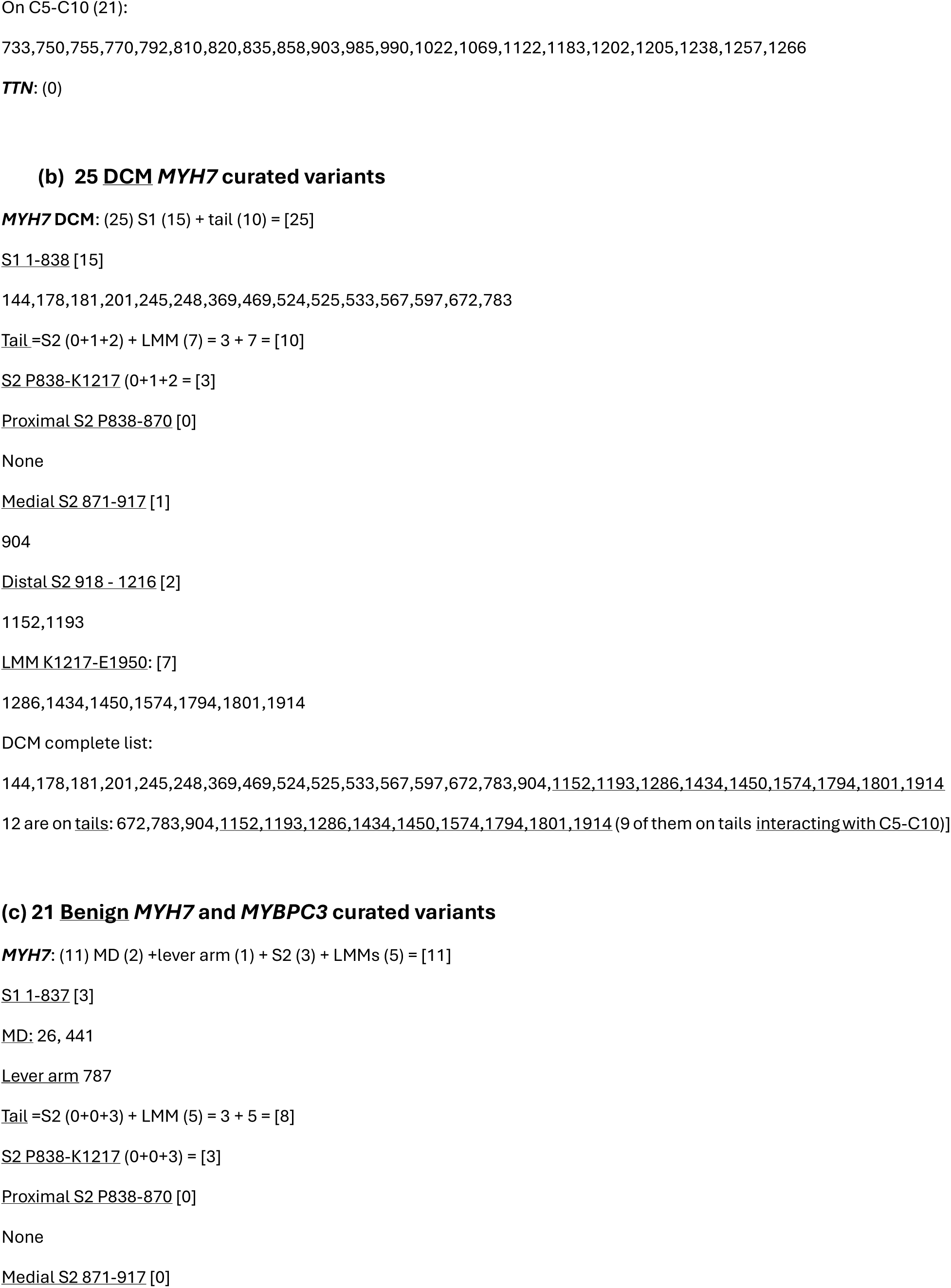
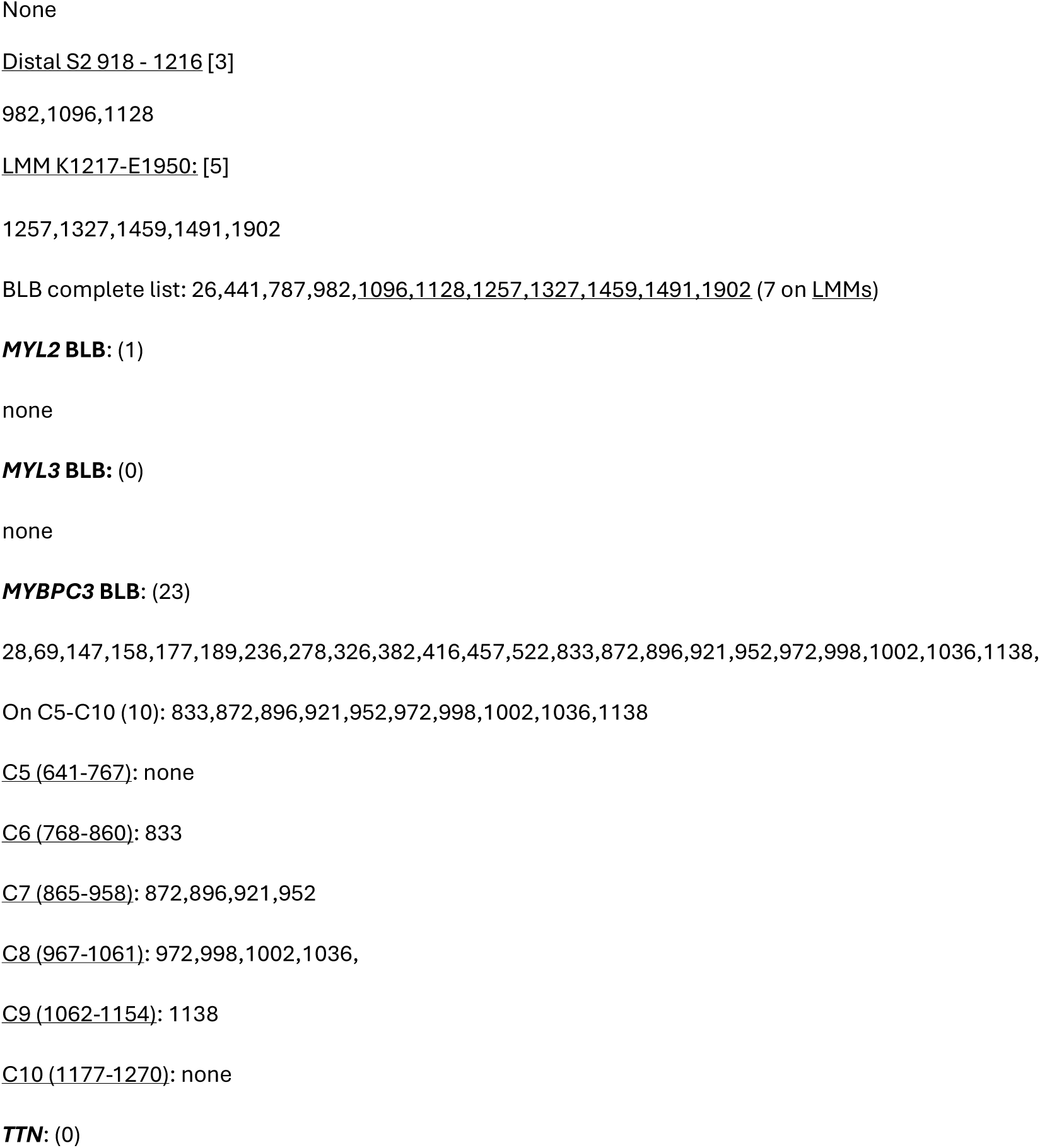
List of HCM, DCM, and benign variants curated from ClinVar.

**Supplementary Table 4.**
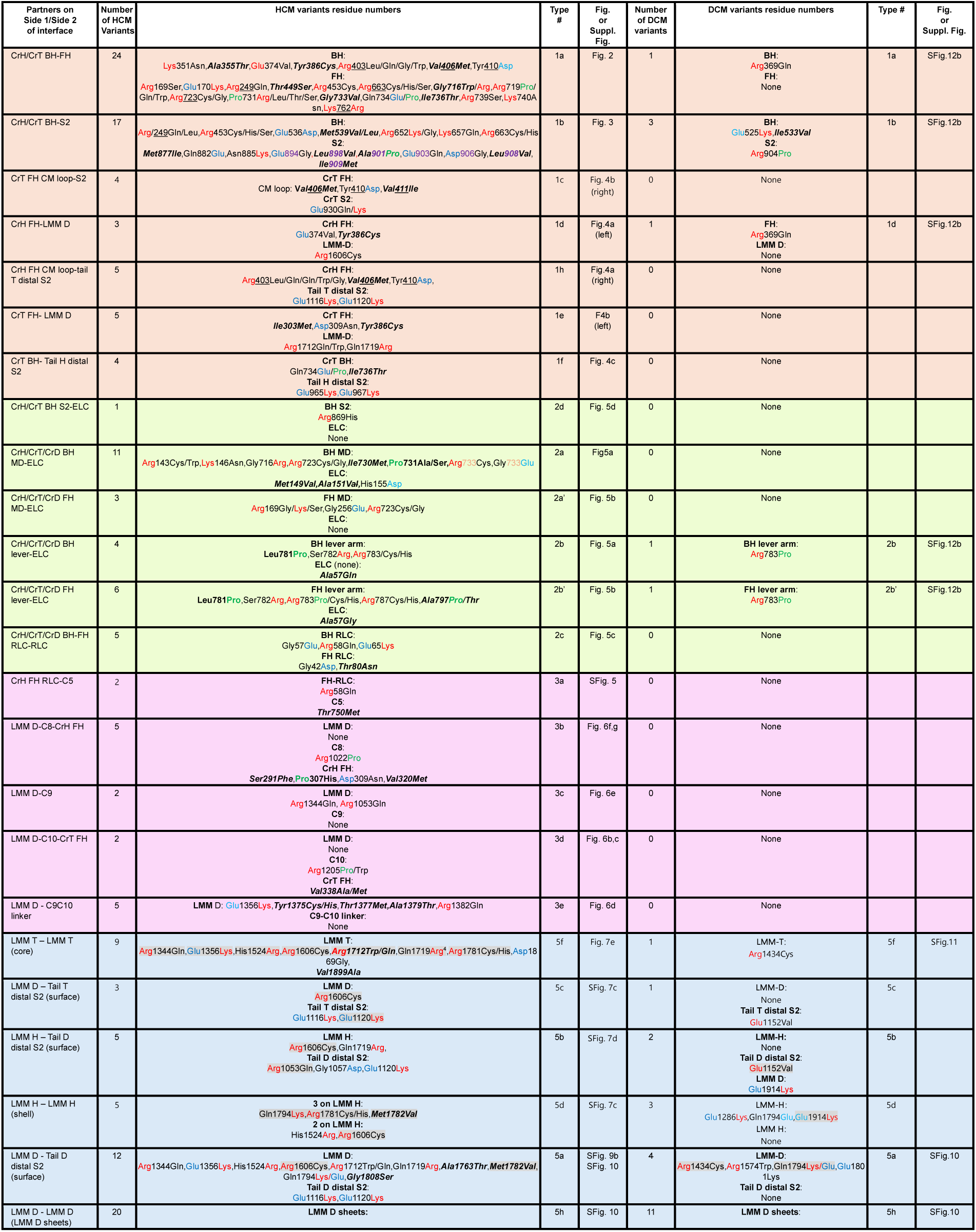
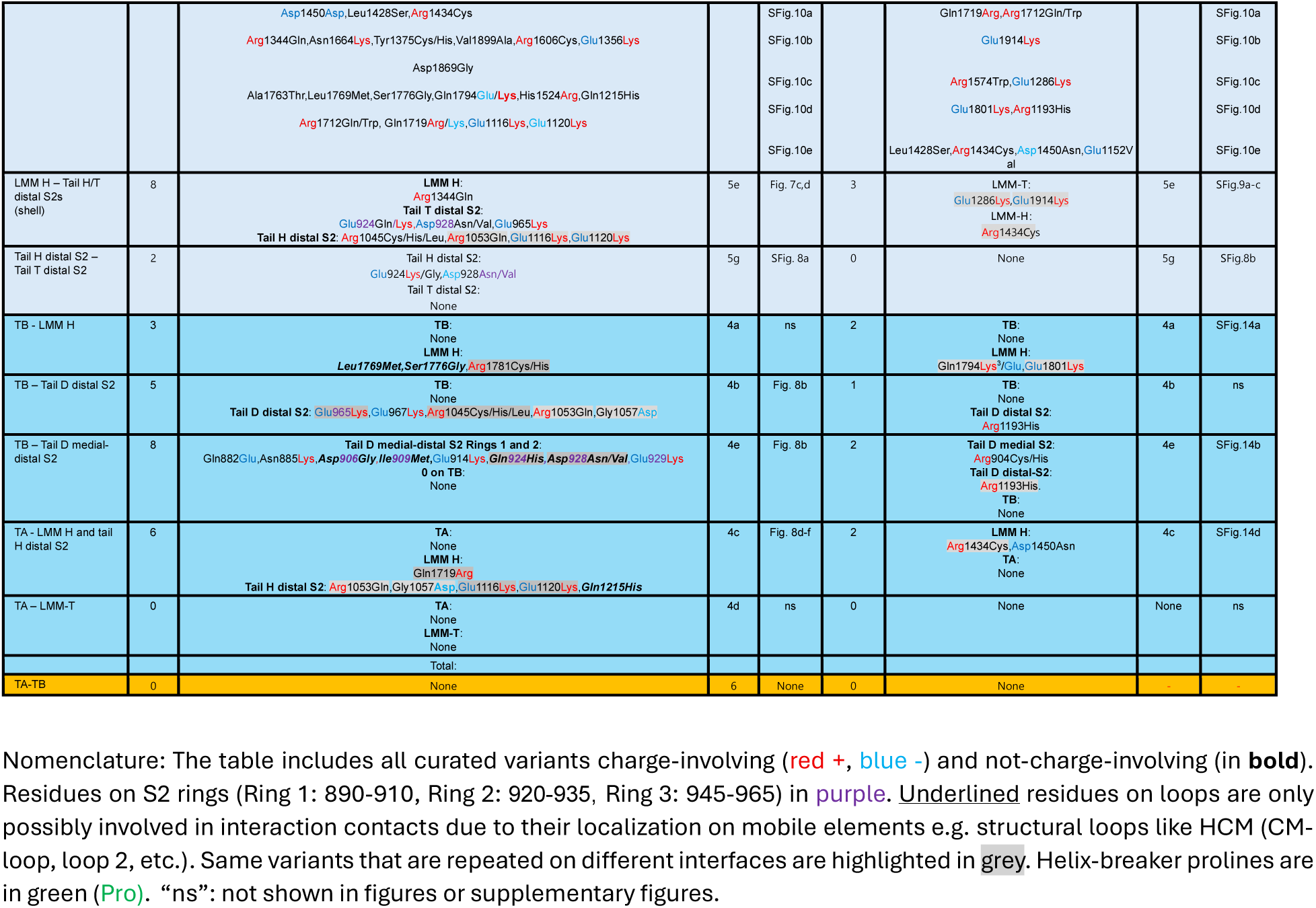
Annotated HCM and DCM variants on the 32 interfaces of the human cardiac thick filament.

**Supplementary Table 5.** Clustering of *MYH7* pathogenic HCM variants annotated in ClinVar *.

|  | BH-FH interface | BH-S2 interface | Proximal S2 | Medial S2 | Rest of tail (distal S2/LMM)<sup>c</sup> |
| --- | --- | --- | --- | --- | --- |
| 1. Total number of residues in each interface or region of interest (ROI) (Supp. Datafile 1, Supp. Table 3) | 61 | 70 | 32 | 47 | 1018 |
| 2. Number of ClinVar pathogenic variants in the interface or ROI <sup>a</sup> (Supp. Table 3) | 24 | 17 | 11 | 12 | 35 |
| 3. Expected number of variants in interface or ROI assuming uniform distribution <sup>b</sup><br>[Row 1/Row 4] x Row 5 | 8.5 | 8.9 | 1.7 | 2.5 | 92 |
| 4. Region used for comparison and its total number of residues (Supp. Table 3) | MHC head/ 838 | MHC S1+S2/ 1217 | MHC tail/ 1097 | MHC tail/ 1097 | MHC whole molecule/ 1935 |
| 5. Total number of variants in region of comparison (Supp. Table 3) | 117 | 154 | 58 | 58 | 175 |
| 6. Clustering factor<br>Row 2/Row 3 <sup>d</sup> | 2.8 | 1.9 | 6.5 | 4.8 | 0.38 |
| 7. P value <sup>e</sup> | 3e-06 | 0.009 | 8e-8 | 5e-6 | 4e-12 (protected) |

\*There are 215 pathogenic variants in ClinVar (Supplementary Datafile 2) that alter 175 of the 1935 residues comprising the MHC
The table shows clustering of many of these variants in specific regions of the MHC: the BH-FH interface, the BH-S2 interface, and different regions of the tail (proximal S2, where it leaves the heads; medial S2, where it crosses the BH; the rest of the tail after it leaves the BH, i.e. distal S2 and LMM).
The degree of clustering in these different regions is determined by comparing the number of variants that would be expected in a location if they were evenly distributed over the molecule (or relevant local region of the molecule) compared with the number found clinically.
<sup>a</sup> The number of residues impacted by pathogenic variants in each interface or ROI, curated from ClinVar
<sup>b</sup> The expected number of variants in the interface or ROI is calculated based on the number of amino acids in the interface or ROI (Row 1), the region used for comparison and its number of amino acids (Row 4), and the total number of variants in that region (Row 5). Sample calculation: Expected variants for BH-FH interface (containing 61 residues): $[61/838] \times 117 = 8.5$ .
<sup>c</sup> Rest of tail = the region beyond the IHM – i.e., the region forming the filament backbone
<sup>d</sup> Factor by which region is enriched in variants compared with uniform distribution
<sup>e</sup> Likelihood that enrichment factor would occur by chance, based on Monte Carlo simulations.

